# Deletion of *Xist* upstream sequences alters TAD interactions and leads to defects in Xist coating and expression

**DOI:** 10.1101/2023.08.14.553118

**Authors:** S Majumdar, LS Bammidi, HC Naik, Avinchal, R Baro, A Kalita, N Sundarraj, GS Bariha, D Notani, S Gayen

## Abstract

The topological organization of the genome plays an important role in regulating gene expression. However, the connection between the two remains poorly understood. X-chromosome inactivation is a unique model system to explore the interlink between topologically associated domains (TADs) and gene expression. TADs are largely lost upon X-inactivation, and the inactive-X gets bipartitely reorganized into two large mega domains. However, the X-inactivation center (XIC) harbors two TADs – at the locus of long non-coding RNA Xist (Xist-TAD) and Tsix (Tsix-TAD). Xist is the master regulator of X-inactivation, which coat the inactive-X and facilitates heterochromatinization. Here, we deleted Xist upstream sequences (∼6 kb) near the Xist TAD’s boundary in extraembryonic endoderm stem cells (XEN), which undergo imprinted X-inactivation. This deletion led to the major rearrangement of TADs and affected the expression of genes located within Xist and Tsix TAD, specially the expression of Xist was upregulated, suggesting TADs are essential for proper transcriptional regulation. On the other hand, Xist-upstream deletion on the inactive-X resulted in dispersal of Xist coating and loss of enrichment of repressive chromatin marks on the inactive-X but no effect on X-linked gene silencing. However, we found that autosomal genes were dysregulated in Xist-upstream deleted cells, probably because of misregulation of genes located in Xist and Tsix-TAD, specially Xist. We conclude that Xist upstream sequences are necessary for proper organization of the TADs at the XIC, maintenance of Xist coating/expression and autosomal gene expression.

## Introduction

Eukaryotic genomes are compacted into a minuscule-sized cell nucleus through different layers of structural organizations such as compartmentalization, formation of topologically associated domains (TADs), chromatin loops etc. Among these, TADs are one of the central structural units which comprise self-interacting domains demarcated by chromatin-associated CCCTC-binding factor (CTCF) and the chromatin loop extruder-Cohesin (Dixon et al., 2012; Fraser et al., 2015; Fudenberg et al., 2017; Nora et al., 2012; Szabo et al., 2019). It is believed that TADs are essential for the proper regulation of genome functions such as transcriptional regulation, chromatin state etc. (Collombet et al., 2023; Ibrahim and Mundlos, 2020; Lupiáñez et al., 2015). However, recent studies exploring the role of TADs in regulating transcriptional gene expression have resulted in mixed outcomes. It has been shown that perturbation of TADs does not always affect gene expression raising the question of how vital TADs are for transcriptional regulation (Despang et al., 2019; Ghavi-Helm et al., 2019; Narendra et al., 2016; Nora et al., 2017; Paliou et al., 2019; Williamson et al., 2019; Zuin et al., 2014).

Here, we have explored the relationship between TAD and transcription using the topological organization of the mouse inactive-X chromosome. X-inactivation is a process by which female mammals compensate the dosage of X-linked gene expression between sexes (Gayen et al., 2015, 2016; Lyon, 1961; Saiba et al., 2018). In mouse, there are two forms of X-inactivation: imprinted and random (Harris et al., 2019; Maclary et al., 2017; Mandal et al., 2020; Sarkar et al., 2015). At the onset of early embryonic development (∼4-cell stage), paternal-X undergoes imprinted X-inactivation and switches to random in embryonic epiblast cells, where either paternal or maternal-X is chosen for the inactivation (Maclary et al., 2014; Naik et al., 2022; Samanta et al., 2022). X-inactivation is orchestrated by a series of regulatory elements encoded from the X-inactivation center (XIC), including the master regulator Xist lncRNA and also its antagonist Tsix (Gayen et al., 2015; Lee and Lu, 1999; Yin et al., 2021). Xist RNA spreads and coats the inactive-X and recruits different chromatin modifiers to enable heterochromatinization of the inactive-X, while Tsix prevents Xist transcription from the active-X to keep it transcriptionally active (Gayen et al., 2015; Maclary et al., 2014). Beyond this, emerging trends suggest that the 3D-organization of the inactive-X is another critical aspect for orchestration of the X-inactivation. Indeed, upon initiation of random X-inactivation in differentiated embryonic stem cells (ESC), the TADs are largely lost from the inactive-X with the concomitant reorganization of the inactive-X into two large mega-domains (Deng et al., 2015; Giorgetti et al., 2016; Loda et al., 2022). Only a few TAD-like structures remain at the inactive-X after it is restructured, with most of them associated with regions of chromatin accessibility and gene expression (Giorgetti et al., 2016). Interestingly, XIC organized into two TADs – Tsix-TAD (comprised of negative regulators of *Xist* like *Tsix*, *Xite*) and Xist-TAD (consisting of *Xist* and its positive regulators *Jpx*, *Ftx*, *Rnf12*), which function in a complete antagonistic fashion with genes in Tsix-TAD getting downregulated and that of Xist-TAD preferentially upregulated from the inactive-X upon initiation of X-inactivation (van Bemmel et al., 2019). Altogether, it is believed that proper topological organization of XIC is important for orchestrating X-inactivation. However, the topological organization of XIC in cells with imprinted X-inactivation remains underexplored. Here, we have explored the TADs and chromatin state of XIC in mouse extra-embryonic endoderm stem cells (XEN), which undergo imprinted X-inactivation. We show that XIC in XEN cells harbour Xist and Tsix TAD, similar to the other cell types with random X-inactivation. Furthermore, how crucial is the proper topological organization of the XIC in maintenance of X-inactivation remains poorly understood. To explore this, we deleted Xist upstream intra-TAD sequences near the TAD boundary in XEN cells. Interestingly, we found that deletion of Xist-upstream sequences destabilizes the contacts of TAD and affect the expression of genes located in Xist and Tsix-TADs, specially Xist, suggesting TADs are important for transcriptional regulation. Furthermore, we found that Xist-upstream sequences result in the dispersal of Xist coating and loss of enrichment of repressive chromatin marks but do not affect X-linked gene silencing. Surprisingly, despite no effect on inactive-X gene silencing, the expression of many autosomal genes was dysregulated in Xist-upstream deleted cells, probably due to the dysregulation of Xist and other TAD-associated gene expression. Altogether, our study reveals that Xist upstream sequences are important for maintaining proper TAD organization, Xist coating/expression, autosomal gene expression and provide significant insight into the role of proper genome organization in regulating transcriptional gene expression.

## Results

### Topological organization of the XIC of imprinted inactive-X in XEN cells is similar to cells with random inactivated-X but have different chromatin state

It is thought that proper topological organization and chromatin states of the XIC are important for orchestration of X-inactivation. Although, TADs are primarily lost upon X-inactivation, XIC contains two TADs: named as Xist and Tsix TAD, as described in different studies involving random X-inactivation (van Bemmel et al., 2019). Tsix-TAD comprised of negative regulators of *Xist* like *Tsix*, *Xite* and Xist-TAD consisting of *Xist* and its positive regulators *Jpx*, *Ftx*, *Rnf12* (Fig. 1A). However, the topological organization of XIC of imprinted inactive-X remains underexplored. To explore this, we have studied the topological organization of the XIC in XEN cells, which undergo imprinted X-inactivation. First, we performed circular chromosome conformation capture sequencing (4C-seq) in XEN cells with a viewpoint (VP 1) at the *Xist* promoter to identify the interacting loci within the XIC. We found major contacts of *Xist* promoter with *Jpx*, *Ftx* and *Slc16a2* loci, suggesting the existence of similar Xist TAD in the XIC of XEN cells. Interestingly, many contacts overlapped with the contacts identified through Capture-C analysis of differentiating female ESC with random X-inactivation using Xist viewpoint (Fig. 1C). Another viewpoint (VP2) analysis near the Xist promoter also showed few contacts within the Xist TAD (Fig. 1C). Taken together, our 4C-seq analysis revealed the existence of Xist TAD in XEN cells. However, we observed contacts of Xist loci with a few loci within the adjacent Tsix-TAD for both of the view-point, suggesting the existence of inter TAD interactions (Fig. 1C). Next, we profiled the enrichment of CTCF insulator protein in XEN cells across the XIC in active-X vs. inactive-X chromosome through allele-specific chromatin immunoprecipitation sequencing (ChIP-seq) analysis (Fig. 1C). XEN cells used for this study is a hybrid cell line derived from two divergent mouse strains *Mus musculus* (mus) and *Mus molassinus* (mol), which allowed us to perform allele-specific analysis (Fig. 1B). Moreover, paternal X is inactivated in XEN cells which helped us to distinguish between active-X vs. inactive-X specific enrichment. Notably, many contact loci identified through 4C analysis showed enrichment of CTCF (Fig. 1C). Furthermore, we demonstrate that CTCF enrichment spanning the XIC in XEN cells is very similar to the enrichment in neural progenitor cells (NPC) with random X-inactivation (Fig. 1C). Taken together, our analysis demonstrates that topological organization of the XIC of imprinted inactive-X in XEN cells is very similar to the random X-inactivation. On the other hand, enrichment of enhancer marks like H3K4me1 and H3K27ac, which regulate chromatin loop and domain formation, showed striking differences between XEN vs. differentiated ESC undergoing random XCI (Fig. 1C). While H3K4me1 and H3K27ac showed strong enrichment on Xist-TAD in XEN cells, it was strongly enriched on Tsix-TAD in differentiated ESC (Fig. 1C). Collectively, we show that while the topology of the XIC is very similar between XEN cells and differentiating ESC, chromatin enrichment is different.

**Figure 1:**
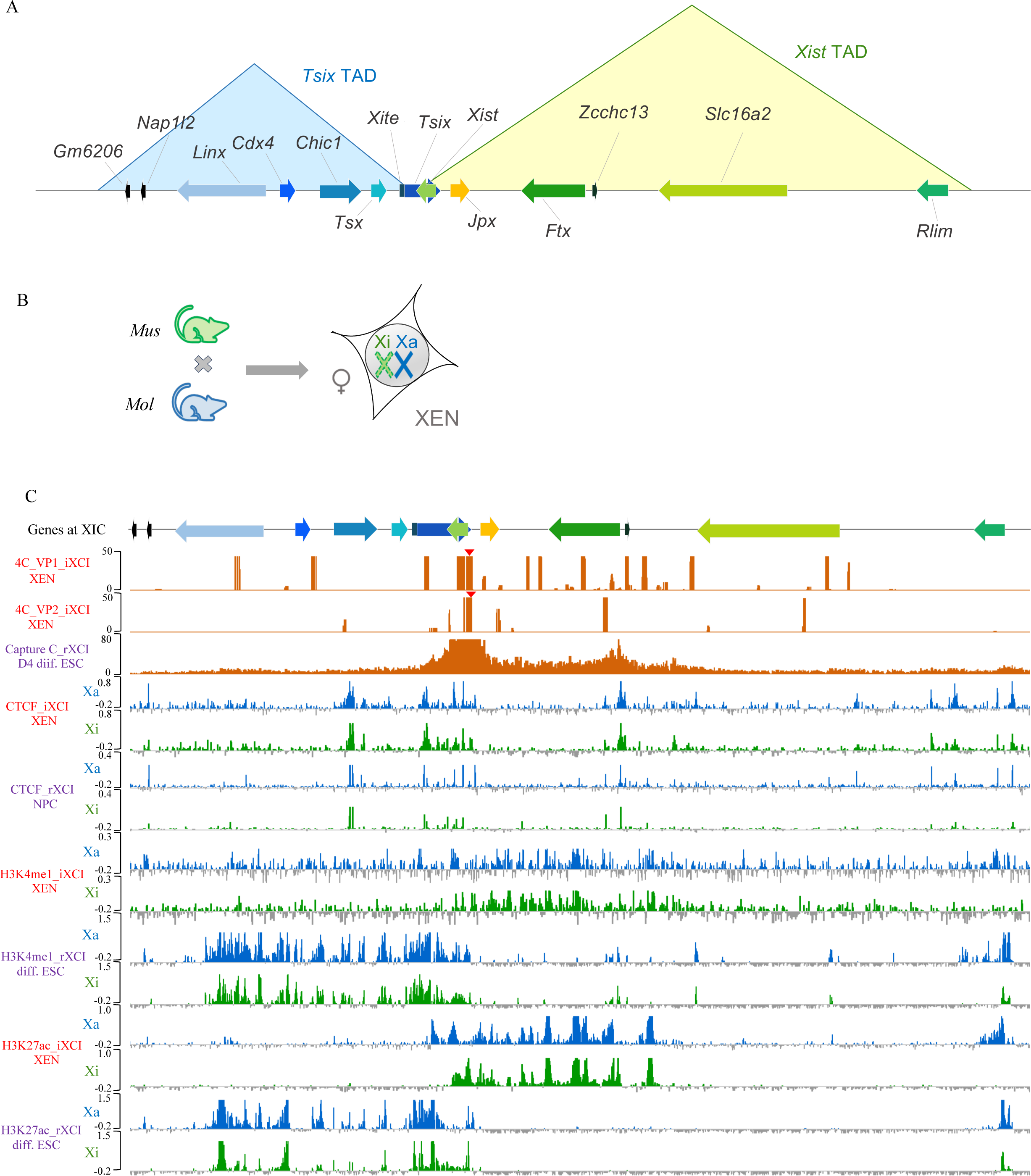
Comparison of topological organization and chromatin features of the XIC between imprinted and random X-inactivation. (A) Schematic representing the linear arrangement of genes across the Xist and Tsix TADs. The Tsix TAD is shaded in blue and the Xist TAD in green. (B) Diagrammatic representation of hybrid XEN cells derived from a cross between *Mus musculus* (mus) and *Mus molossinus* (mol) mice. The X chromosome derived from *mus* is the inactive-X. (C) Topological organization and chromatin features of the XIC in XEN cells (imprinted) and cells undergoing random X-inactivation. Top: 4C interaction profile of Xist VP1 and VP2 in XEN cells and Capture-C profile of Xist in day 4 differentiating ESCs undergoing random X-inactivation. Bottom: ChIP-seq profiles of the XIC in active-X (blue) and inactive-X (green) for CTCF, H3K4me1 and H3K27ac marks in XEN cells vs. NPCs or differentiating ESCs undergoing random X-inactivation. iXCI: imprinted X-chromosome inactivation; rXCI: random X-chromosome inactivation

### Deletion of *Xist* upstream sequences leads to alterations of interactions within TAD and across the X-chromosome

Next, we investigated the importance of proper topological organization of the XIC in the maintenance of imprinted XCI in XEN cells. For this, we deleted *Xist*-upstream sequences (∼6kb) lying within the Xist TAD near the TAD boundary in XEN cells keeping intact the promoter of the Xist (Fig 2A). Moreover, this region does not have major CTCF peaks or enrichment of H3K4me1 and H3k27ac marks (Fig S1A). We generated two heterozygous XEN cell lines with paternal deletion (D1 ΔXp.1, D1 ΔXp.2) and three heterozygous clones with maternal allele deletion (D1 ΔXm.1, D1 ΔXm.2, D1 ΔXm.3) using CRISPR/Cas9 (Fig. 2B). The paternal or maternal deletions were verified through SNP-based Sanger-sequencing (Fig. S1B and S1C). We also generated empty-vector transfected clonal lines (WT EV1, WT EV2 and WT EV3) for our experiments.

**Figure 2:**
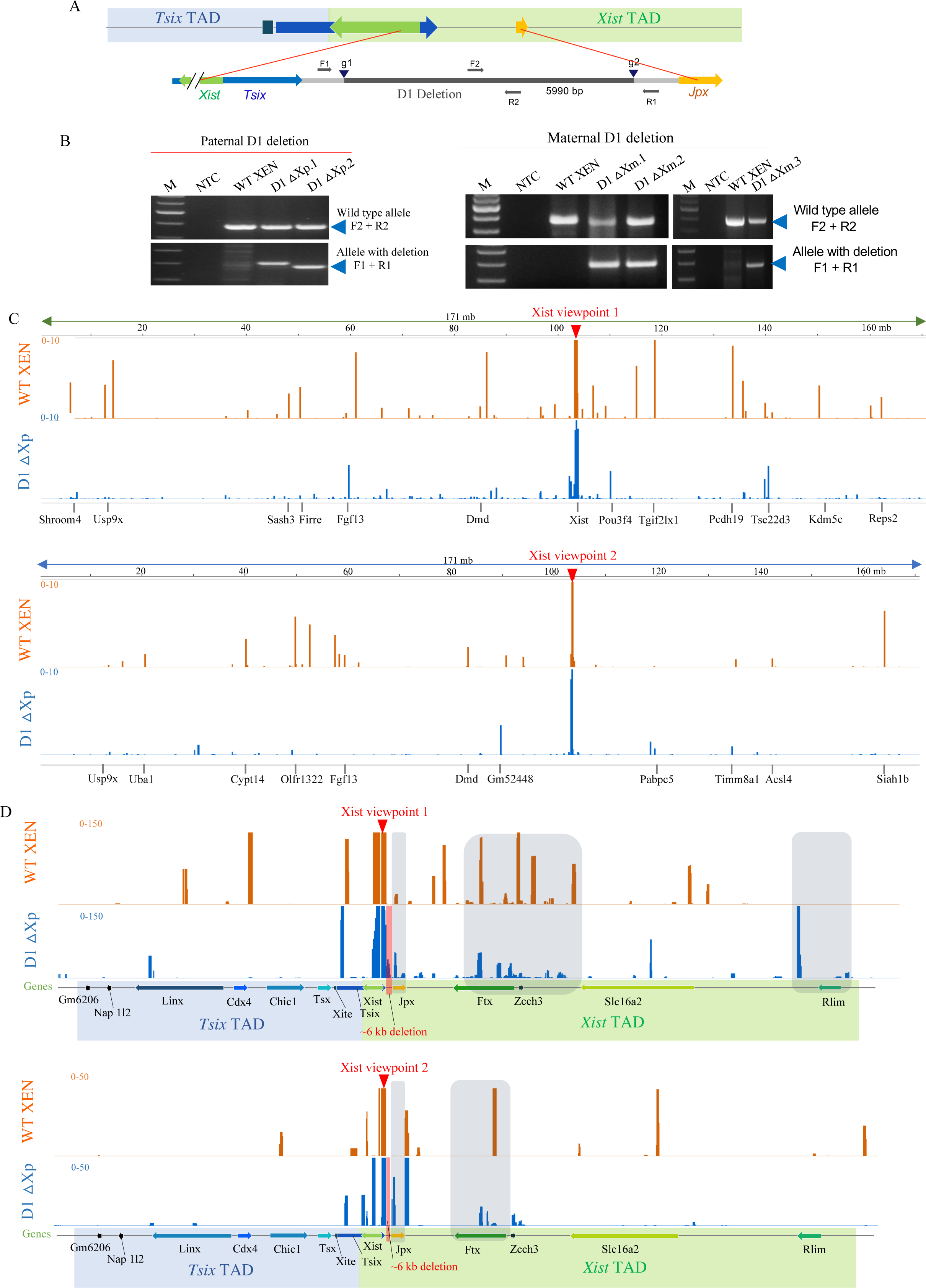
Generation of *Xist* upstream sequence deleted XEN cells clones and profiling of interactions of *Xist* locus upon *Xist* upstream deletion. (A) Schematic showing the region intermediate to *Xist* and *Jpx* that is targeted for deletion in XEN cells. sgRNA-binding sites: g1 and g2. PCR primers F2 and R2, and F1 and R1 shown as grey arrows used for amplification of the wild-type allele and the allele with deletion respectively. (B) PCR-based identification of clones having the deletion of the *Xist* upstream region (D1) on the paternal-allele (left) and maternal-allele (right). Arrowheads represent the position of the expected amplicon. Primer pairs used for each deletion are shown on the right. NTC stands for no template control and M for Marker. (C) 4C interaction profile for *Xist* promoter region with VP1 and VP2 (marked as red arrowhead) across the entire X-chromosome in wildtype and D1 ΔXp.1. (D) 4C interaction profile of *Xist* promoter region within the XIC. The Xist and the Tsix TADs have been shaded as green and blue in the XIC region. The deleted region have been highlighted in red in the D1 ΔXp 4C profiles. Differential interactions between the WT and D1 ΔXp have been shaded in grey.

First, we investigated, whether the deletion of *Xist* upstream sequences on the inactive-X affects the TAD organization. To explore this, we performed 4C-seq on a paternally deleted line considering two viewpoints (VP1 and VP2) on Xist (Fig. 2). We found that Xist upstream sequence deletion in D1 ΔXp.1 clone leads to the loss of interaction with the Xist locus chromosome-wide (Fig 2C). Moreover, in the case of VP1, there were significant alterations of contacts within Xist TAD and neighbouring Tsix-TAD (Fig. 2D, Fig. S2A). Notably, the interaction between Xist and Ftx was drastically reduced with the gain of interactions between Xist and Rnf12 (Fig. 2D). Similarly, for VP2, there was loss of contacts within Xist and Tsix TAD (Fig. 2E Fig. S2B). On the other hand, there were changes of inter-chromosomal contacts upon deletion of *Xist* upstream sequences (Fig. S2C). Taken together, our analysis suggested that the Xist upstream sequences are essential for maintaining TAD contacts in XEN cells.

### Deletion of *Xist* upstream sequence leads to abnormal Xist coating and upregulation of Xist expression

Next, we explored the effect of the *Xist* upstream sequence deletion and associated TADs alterations on Xist coating and expression. To investigate this, we performed RNA fluorescence in-situ hybridization (RNA-FISH) analysis for Xist in WT EV, D1 ΔXp and D1 ΔXm XEN cells. Interestingly, we found that clones (D1 ΔXp) having the deletion on their paternal-allele showed highly dispersed Xist clouds (∼31-42%) (Fig. 3A and 3B). However, the maternally deleted (D1 ΔXm) or the empty-vector transfected clones (WT EV1, WT EV2 and WT EV3) did not show such kind of Xist dispersal. Additionally, a significant percentage of cells in paternally deleted clones (D1 ΔXp) showed faint Xist clouds (Fig. 3B). Next, we compared the area and the sphericity of the dispersed Xist of D1 ΔXp lines with the compact Xist of WT EV clones and found that while the area of the dispersed Xist clouds of the paternally deleted clones (D1 ΔXp) increased significantly, the sphericity of the clouds became significantly lower (Fig. 3C). Collectively, our analysis suggested that the *Xist* upstream region is essential for proper Xist coating in XEN cells.

**Figure 3:**
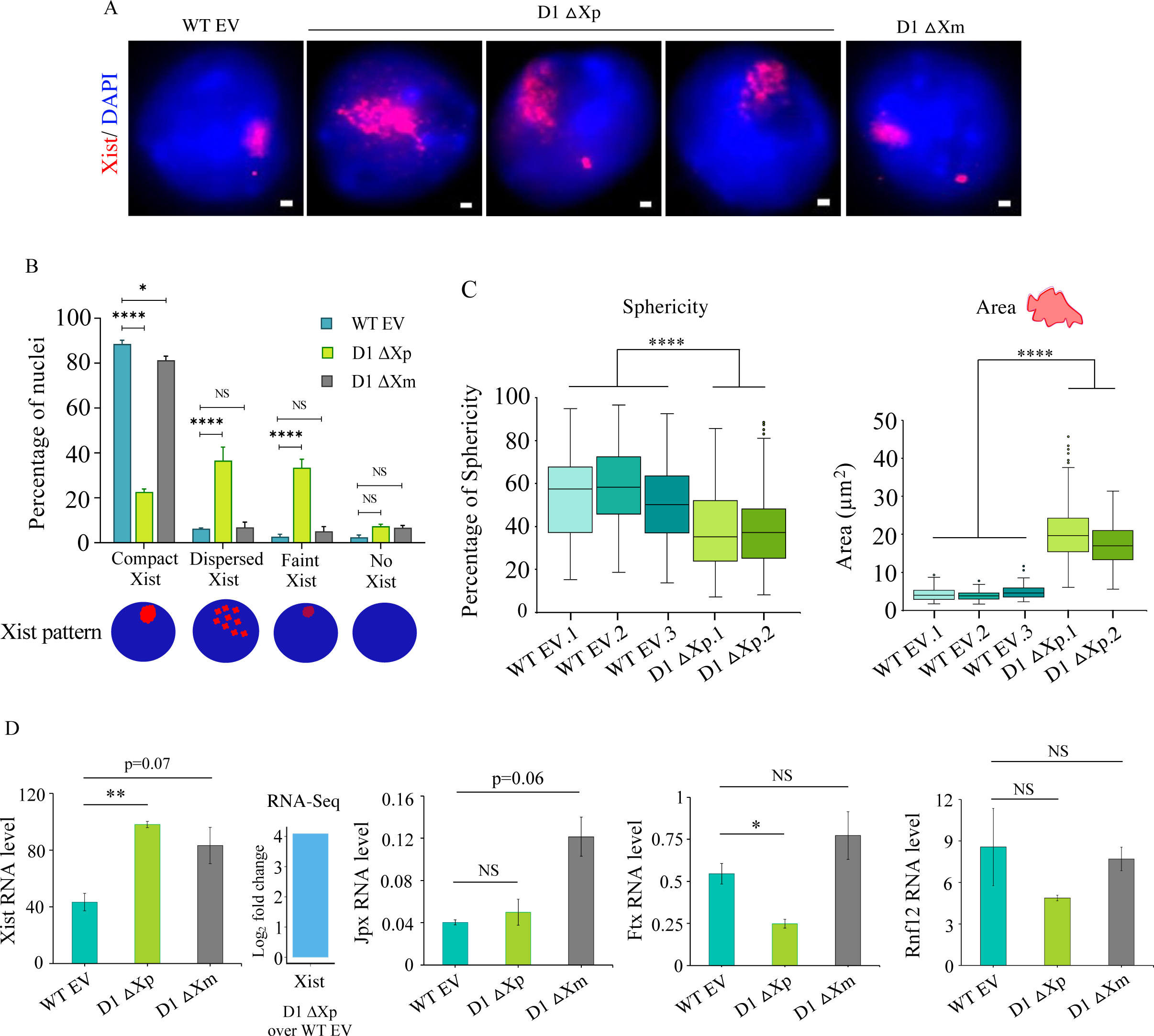
*Xist* upstream deletion leads to Xist dispersal and *Xist* upregulation. (A) Representative RNA-FISH images of Xist/Tsix (red) in wild-type empty-vector transfected (WT EV) and heterozygous clones of both paternally (D1 ΔXp) and maternally (D1 ΔXm) deleted clones. DAPI (blue) counterstains the nucleus. Scale bars: 1µm. (B) Plot representing the quantification of the Xist RNA-FISH signals in WT EV and heterozygous (D1 ΔXp and D1 ΔXm) clones. Bottom: schematic representing different types of Xist pattern. n=3 biological replicates in each of WT EV and D1 ΔXm and 2 for D1 ΔXp. *P* values (two-way Annova) \*\*\*\**P*≤0.0001, \**P*≤0.05 and NS implies non-significant. (C) Boxplots representing the quantification of sphericity and the area occupied by Xist clouds in WT EV clones and clones with paternal D1 deletion. *P* values (Mann-Whitney test) as compared to wildtype; \*\*\*\**P*≤0.0001. n= 113, 100, 110, 100, 103, 95. (D) Analysis of *Xist*, *Jpx*, *Ftx* and *Rnf12* mRNA expression levels by qRT-PCR in WT EV and D1 deleted clones. TBP is used as the normalizing control. n=3 biological replicates for each of WT EV and D1 ΔXm and 2 for D1 ΔXp was used. Error bars represent the standard error of mean. *P* values (Two-tailed t –test): \**P*≤0.05, \*\**P*Δ0.01. log_2_ fold change for Xist in D1 ΔXp over WT EV clones as seen through RNA-sequencing has been represented beside the qRT-PCR plot for Xist.

We then performed quantitative RT-PCR analysis to explore if *Xist* upstream deletion affects the expression of *Xist,* including other *Xist* regulators (*Jpx, Ftx, Rnf12*) located within the same Xist TAD. We found that the expression of Xist in paternally deleted clonal lines (D1 ΔXp) is significantly increased compared to the WT (Fig. 3D). RNA-seq analysis confirmed the same (Fig. 3D). On the other hand, we did not notice any significant alteration in expression of *Jpx* and *Ftx* expression between WT vs. paternal deletion clones (D1 ΔXp) (Fig. 3D, Fig. S3). In case of *Rnf12*, RNA-seq analysis showed a slight increase in expression in D1 ΔXp, however, qRT-PCR analysis showed little decreased expression. This disimilarity could be due to the different isoforms of *Rnf12*, which may not have covered in qRT-PCR expression analysis. Additionally, the expression of *Chic1*, a gene located in the adjacent Tsix TAD, increased in D1 ΔXp clones (Fig. S3). Taken together, our analysis suggested that perturbation of the Xist upstream region on the inactive-X alters the expression of genes within Xist TAD as well as in neighbouring TAD. On the other hand, in maternally deleted clonal lines (D1 ΔXm), there was a slight increase in expression for *Xist* and *Jpx*, whereas the expression of *Rnf12* or *Ftx* remains unaffected (Fig. 3D).

### Segmental deletion of the *Xist* upstream region revealed that region near *Jpx* is essential for proper Xist coating

Next, to further explore the link between *Xist* upstream region and proper Xist coating, we generated XEN cell lines with two segmental deletions (D2: near to *Xist* and D3: near to *Jpx*) of the *Xist* upstream region using CRISPR/Cas9 (Fig. 4A). We generated heterozygous clones having deletion from either paternal or maternal allele for each of the D2 and D3 deletions. We validated heterozygous deletions through PCR followed by Sanger-sequencing (Fig. 4B and 4C, Fig. S4A,B,C and D). We also obtained homozygous clones for the D3 deletion (D3 ΔXpΔXm.1 and D3 ΔXpΔXm.2). Next, to explore the status of Xist coating, we performed RNA-FISH experiments for Xist in the D2 and D3 deleted XEN cell clones. Interestingly, we found that deletion of the region near to *Jpx* (D3) on the inactive-X allele leads to the dispersal of Xist coating to an almost similar extent (∼32-40% nuclei) as it was observed in the paternally deleted lines of the D1 deletion (Fig. 4E and 4F). In consistence, the dispersed Xist clouds of D3 homozygous and paternally deleted clones exhibited greater area and decreased sphericity compared to the compact Xist clouds of WT clones (Fig 4G). In converse, D3 deletion from maternal/active-X allele did not lead to such defects on Xist coating (Fig. 4E). However, homozygous deletion of D3 led to the dispersal of Xist coating suggesting the *cis*-effect of D3 deletion towards maintaining proper Xist coating (Fig. 4E). On the other hand, the D2 deletion from paternal/inactive-X or the maternal/active-X allele did not show such defect in Xist coating (Fig. 4D). Collectively, our analysis suggested that the *Xist* upstream sequence near to *Jpx* is essential for proper Xist coating.

**Figure 4:**
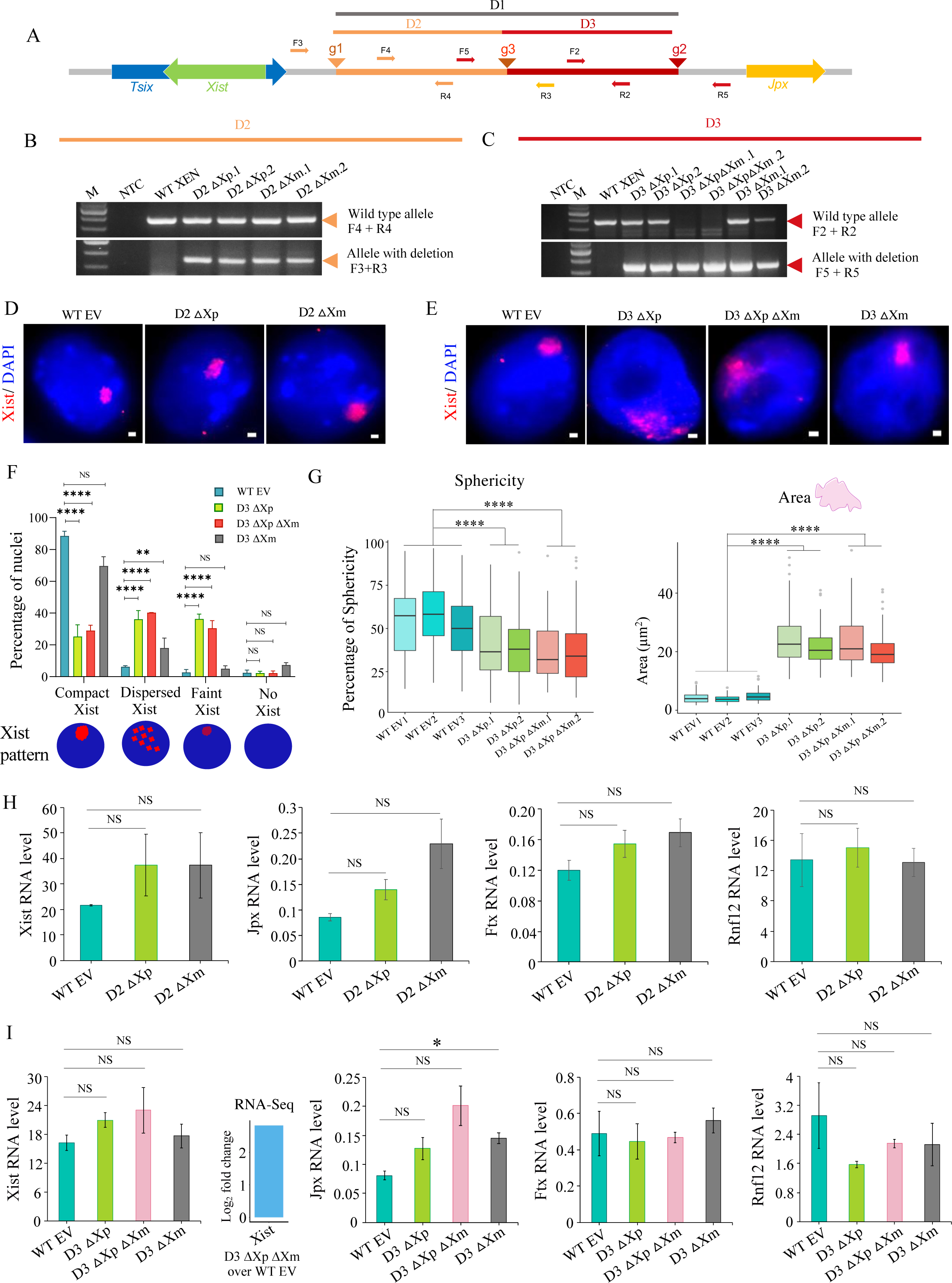
Segmental deletions of the *Xist* upstream sequence leads to transcriptional changes of genes within Xist/Tsix TADs and dispersal of Xist coating. (A) Schematic showing the segmental deletions (D2 and D3) regions upstream to *Xist*. Guide RNA target sites are represented as orange arrowheads (g1 and g3) for the smaller D2 deletion and as red arrowheads (g3 and g2) for the D3 deletion. Primer pairs are shown as orange and red arrows, F3 and R3 amplify the deleted allele and F4 and R4 amplify the wildtype allele of the D2 deletion. F2 and R2 amplify the undeleted allele and F5 and R5 amplify the deleted allele of the D3 deletion. (B) PCR based identification of D2 paternally (D2 ΔXp.1 and D2 ΔXp.2) and maternally deleted clones (D2 ΔXm.1 and D2 ΔXm.2). Orange arrows mark the position of the expected amplicons. M denotes marker and NTC: no template control. (C) PCR for the paternally (D3 ΔXp.1 and D3 ΔXp.2), homozygous (D3 ΔXp ΔXm.1 and D3 ΔXp ΔXm.2) and maternally (D3 ΔXm.1 and D3 ΔXm.2) deleted lines of the D3 deletion. Red arrowheads mark the position of the amplicons. The primers used for each PCR are indicated on the right. (D) RNA-FISH for WT EV and D2 deletion clones using double-stranded Xist/Tsix-Cy3 (red) probe. Nuclei are counterstained with DAPI (blue). Scale bars: 1µm. (E) RNA-FISH for WT EV, D3 ΔXp, ΔXpΔXm and ΔXm clones. Xist and Tsix signals are in red. Scale bars: 1µm. (F) Quantification of the pattern of different Xist coating in the D3 deleted clones. n= 3 for WT EV clones and 2 for each of D3 ΔXp, D3 ΔXpΔXm and D3 ΔXm clones. *P* values (two-way Annova) \*\*\*\**P*≤0.0001, \*\**P*≤0.01. (G) Quantification of the sphericity and area occupied by the Xist clouds in the D3 deleted (homozygous and paternally deleted) clonal lines. *P* values (Mann-Whitney test) as compared to WT EV clones. n=113, 100, 110, 99, 102, 98, 97. (H) and (I) qRT-PCR for analysing *Xist*, *Jpx*, *Ftx* and *Rnf12* expression levels normalized to *Tbp* for the D2 and D3 deleted lines respectively. *P* values (Two-tailed t –test): \**P*≤0.05. log2 fold change of *Xist* expression in D3 ΔXpΔXm lines as compared to WT EV clones as observed through RNA-sequencing have been represented at the right of the Xist qRT-PCR plot.

Next, we explored the impact of D2 or D3 deletion on the expression of genes (*Xist*, *Jpx*, *Ftx* and *Rnf12*) located within Xist TAD. We found that both deletions (D2 and D3) from the paternal/inactive-X led to increase expression of *Xist* and *Jpx* (Fig. 4H and 4I). The homozygous D3 deletion also showed the same expression profile for *Xist* and *Jpx* (Fig. 4I, Fig. S4E). Other genes of the Xist TAD, like *Ftx* and *Rnf12* did not show any considerable changes in their expression in any of the D2 or the D3 clonally deleted lines (Fig. 4H and 4I). On the other hand, expression of *Chic1*, a gene located within the Tsix TAD, was increased in D3 homozygous XEN cells (Fig. S4E). Altogether, segmental deletions (D2 and D3) of the *Xist* upstream sequence affected TAD-related gene expression.

### Xist dispersal leads to loss of repressive chromatin marks but no effect on the nucleolar association of the inactive-X

After initiation of X-inactivation, many repressive chromatin marks such as H3k27me3, H2AK119ub, H4K20me, Macro H2A etc., accumulate on the inactive-X, which plays an important role in the maintenance of gene silencing. We investigated whether the dispersal of Xist coating leads to any defects or loss of enrichment of these different repressive chromatin marks on the inactive-X in XEN cells. To explore this, we performed immunofluorescence (IF) for these different repressive marks, followed by RNA-FISH for Xist in WT, D1 and D3 XEN cells. As expected, we found all these marks are robustly enriched on the inactive-X in WT XEN cells (Fig. 5). Strikingly, we observed that while ∼90% of WT XEN cells nuclei have robust enrichment of H3K27me3, there was a loss of enrichment of H3K27me3 in nuclei having dispersed Xist coating in D1 ΔXp, D3 ΔXp and D3 ΔXpΔXm XEN cells (Fig. 5A). However, compact Xist coated nuclei in these cells maintained robust enrichment of H3K27me3 (Fig. 5A). Altogether, our data suggested that dispersal of Xist coating in D1 ΔXp, D3 ΔXp and D3 ΔXpΔXm XEN cells leads to loss of H3k27me3 enrichment. Interestingly, H2AK119ub was still robustly enriched in Xist dispersed nuclei of D1 ΔXp, D3 ΔXp and D3 ΔXpΔXm XEN cells similar to the compacted Xist nuclei (Fig. 5B). On the other hand, enrichment of H4K20me1 was lost in ∼90% of nuclei with dispersed Xist coating in D1 ΔXp, D3 ΔXp and D3 ΔXpΔXm XEN cells (Fig. 5C). Additionally, while cells having compact Xist clouds retained strong and localized enrichment of MacroH2A.1, cells having dispersed Xist clouds possessed only impression of faint and diffused MacroH2A,1 implying that proper Xist coating on the Xi is critical for the association of MacroH2A.1 (Fig. 5D). Taken together, our analysis suggested that Xist dispersal in D1 ΔXp, D3 ΔXp and D3 ΔXpΔXm XEN cells leads to loss of H3k27me3, H4k20me1 and macroH2A enrichment on the inactive-X, however, H2AK119ub enrichment remains maintained robustly.

**Figure 5:**
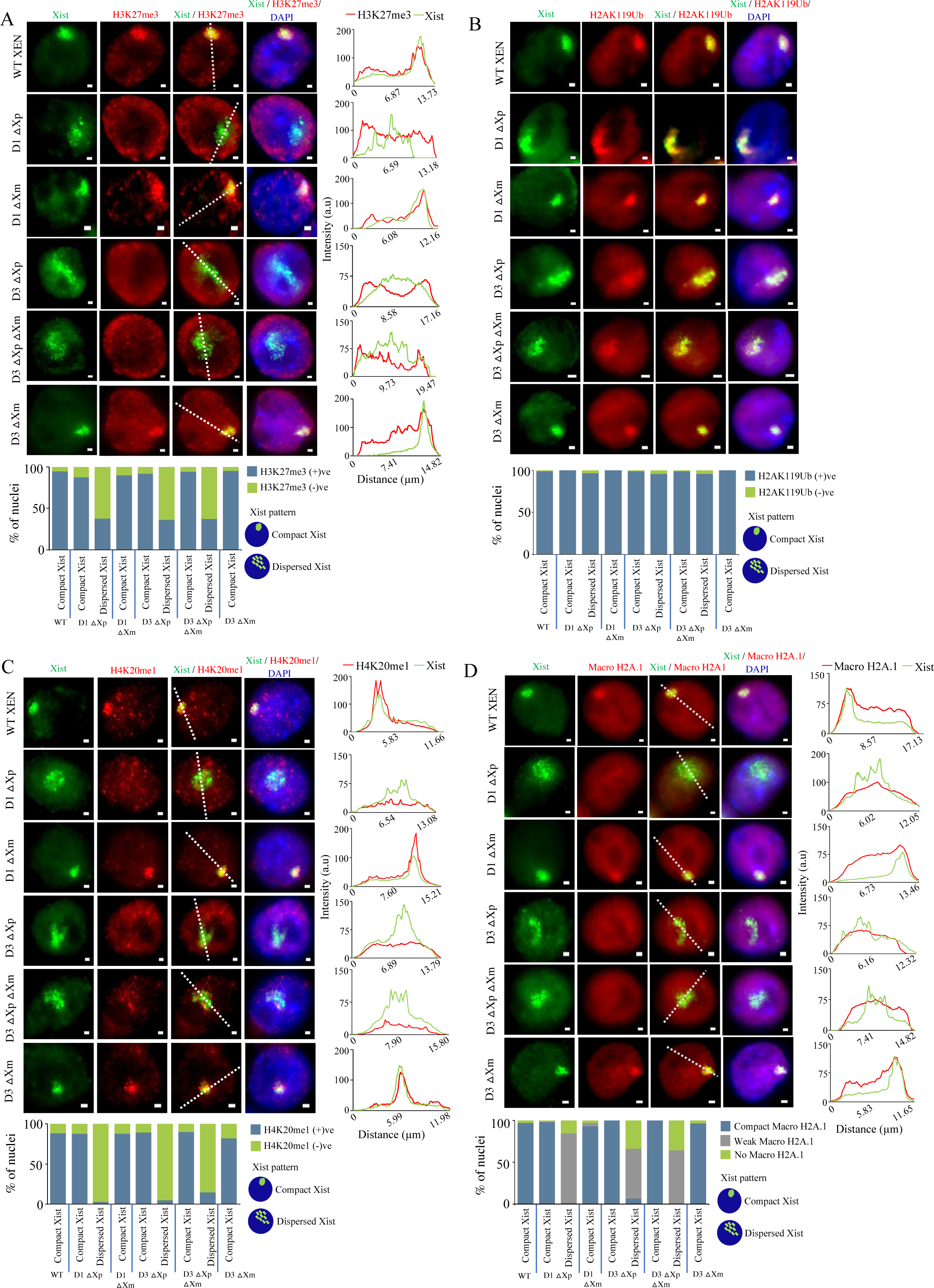
Xist dispersal is associated with the loss of enrichment of repressive chromatin marks on the inactive-X. (A) Representative images of IF-RNA FISH of H3K27me3 (red) and Xist (green) for WT XEN, D1 ΔXp, D1 ΔXm, D3 ΔXp, D3 ΔXp ΔXm, D3 ΔXm lines. Right: corresponding fluorescent intensity profiles for Xist and H3K27me3 along the white line drawn across the Xist cloud. Bottom: quantification of the enrichment of H3K27me3 on the inactive-X in compact and dispersed Xist nuclei. n= 129, 108, 66, 96, 112, 105, 132, 100, 145 (for each data point from left to right). (B) Representative IF-RNA FISH images for H2AK119Ub (red) and Xist (green) for D1 and D3 deleted lines. Bottom: quantification of the the enrichment of H2AK119Ub on the inactive-X in compact and dispersed Xist nuclei. n= 120, 123, 128, 126, 108, 146, 135, 127, 116 (for each data point from left to right). (C) IF-RNA FISH images with corresponding intensity profiles for H4K20me1 (red) and Xist (green). Bottom: quantification for enrichment of H4K20me1 in compact and dispersed Xist nuclei. n= 137, 153, 135, 155, 131, 141, 133, 129, 143 (for each data point from left to right). (D) Macro H2A.1 (red) and Xist (green) IF-RNA FISH representative images with their corresponding intensity profiles are shown. Bottom: quantification of compact, weak and no Macro H2A.1 signals in nuclei having compact and dispersed Xist clouds in the D1 and D3 deleted lines. n= 120, 158, 137, 138, 116, 106, 110, 109, 135 (for each data point from left to right).

Next, we explored whether the dispersal of Xist coating and loss of different repressive chromatin marks leads to the perturbation of nucleolar association of the inactive-X. The inactive-X is known to primarily occupy two regions of the nucleus-one in association with the nucleolus and the other in close proximity to the nuclear membrane (Zhang et al., 2007). The association of the inactive-X to the nucleolus primarily occurs during the maintenance phase of X-inactivation and it is believed that nucleolar association helps to maintain the heterochromatin state of the inactive-X. It is thought that nucleolar association of the inactive-X is primarily facilitated by the Xist RNA and upon loss of Xist, the inactive-X has been found to dissociate from the nucleolus (Kelsey et al., 2015; Zhang et al., 2007). Therefore, to test whether the inactive-X loses its nucleolar association upon Xist dispersal, we performed IF staining for fibrillarin (a protein that is a part of the fibrillar matrix of the nucleolus) to mark the nucleolus followed by RNA-FISH for Xist (Fig. 6A). The Xist clouds were quantified by categorizing its signals to be associated with only the nucleolus, or to the nuclear membrane, or to both the nucleolus and the nuclear membrane or neither of them (Fig. 6A). We found that the dispersed Xist clouds of the D1 ΔXp, D3 ΔXp and D3 ΔXpΔXm XEN cells were similarly associated to the nucleolus as the compact Xist clouds of the wild-type or the maternally deleted lines (Fig. 6A). Taken together, our analysis suggested that inactive-X remain associated with the nucleolus despite of Xist dispersal and loss of repressive marks.

**Figure 6:**
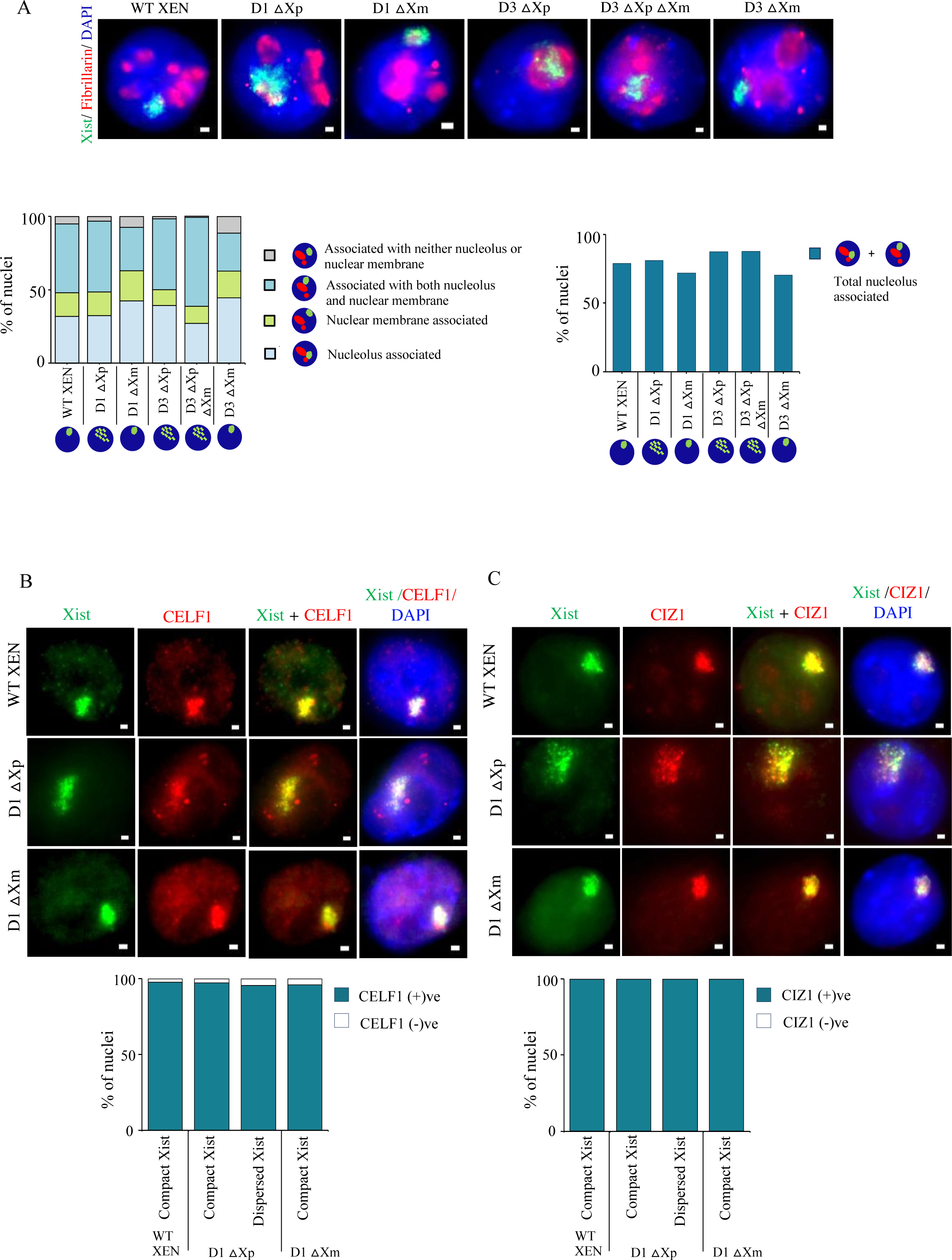
Xist dispersed nuclei retains association with the nucleolus and do not show alteration in CELF1 and CIZ1 enrichment. (A) Top: Representative images of IF-RNA FISH for Fibrillarin (red) and Xist (green) in WT, D1 and D3 deleted lines. Bottom: quantification of the association between Xist cloud and Fibrillarin signals in nuclei with compact and dispersed Xist clouds in the D1 and D3 deleted lines. n= 119, 157, 146, 120, 122, 148 (for each data point from left to right). (B) Images representing IF-RNA-FISH for CIZ1 (red) and Xist (green) in WT and D1 deleted lines. Bottom: quantification of enrichment of CIZ1 on the inactive-X in compact vs. dispersed Xist nuclei. n= 128, 111, 120, 114 (for each data point from left to right). (C) Representative images for IF-RNA-FISH for CELF1(red) and Xist (green) in WT and D1 deleted lines. Bottom: quantification of enrichment of CELF1 on the inactive-X in compact vs. dispersed Xist nuclei. n= 246, 110, 96, 101 (for each data point from left to right).

### Xist tethering proteins remain enriched on the inactive-X of the Xist dispersed nuclei

Next, we wanted to find out if the dispersal of Xist coating is because of the loss of proper enrichment of Xist tethering proteins, such as CELF1, CIZ1 etc., upon Xist upstream deletion / TAD alterations. CELF1 has been identified as a key protein that forms a heterochromatic assembly that maintains proper Xist localization and its loss results in Xist disassociation from the inactive-X (Pandya-Jones et al., 2020). We first looked at the CELF1 enrichment in the D1 and D3 deletion lines that displayed Xist dispersal using immunofluorescence staining for CELF1 coupled with Xist RNA-FISH, but did not find any noticeable defect in the CELF1 enrichment in such nuclei (Fig. 6B). This implied that Xist dispersal was not a consequence of alteration in CELF1 enrichment levels. Next, we looked into the enrichment of another tethering protein CIZ1 that establishes very strong interaction with the Xist RNA and keeps it stably associated with the inactive-X (Ridings-Figueroa et al., 2017). We performed similar experiments for the CIZ1 protein and found that CIZ1 also remained associated with dispersed Xist clouds in paternally deleted D1, D3 and homozygous D3 lines (Fig. 6C). This re-emphasized its strong binding to Xist RNA, which preserved their association even under Xist dispersed conditions. Altogether, our analysis suggested that the dispersal of Xist coating was not associated with the loss of important tethering proteins such as CELF1, CIZ1 etc.

### Xist upstream deletion does not perturb X-linked gene silencing but affects autosomal gene expression

As we observed loss of enrichment of several repressive chromatin marks (H3K27me3, H4K20me1and MacroH2A) on the inactive-X of dispersed Xist nuclei, it intrigued us to explore the status of X-linked gene silencing in inactive-X chromosome. For this, we performed a bulk-RNA sequencing for the WT EV, D1 ΔXp, and D3 ΔXpΔXm XEN cells. As described above, XEN cells used for this study is derived from two divergent mouse strains *Mus musculus* (mus) and *Mus molassinus* (mol), which allowed us to perform allele-specific analysis (Fig. 1B). Moreover, paternal-X (mus) is inactivated in XEN cells which helped us to distinguish between active-X vs. inactive-X specific gene expression through allele-specific analysis of RNA-seq data. As expected, in WT EV clones, X-linked genes showed expression from maternal active-X (X^mol^) (Fig. 7A). Interestingly, D1 ΔXp, and D3 ΔXpΔXm XEN cells as well, most of the X-linked genes expression originated from the maternal active-X (X^mol^) only, suggesting inactive-X still maintains X-linked gene silencing in these cells despite the loss of repressive chromatin marks and Xist dispersal (Fig. 7A). Since RNA-sequencing gave us insight only about the bulk expression profile of the X-linked genes, we wanted to check whether there is X-linked gene reactivation at the single-cell level through RNA-FISH. We performed RNA-FISH for three X-linked genes, namely: *Rnf12*, *Atrx*, *Pg*k1 coupled with *Xist* (Fig. 7B). As expected, WT-XEN cells showed monoallelic expression for all three X-linked genes. Interestingly, dispersed Xist coated nuclei in D1 ΔXp, and D3 ΔXpΔXm XEN cells maintained monoallelic expression similar to the compact Xist coated nuclei (Fig. 7B) Taken together, our analysis suggested that there was no reactivation of X-linked genes from the inactive-X upon Xist dispersal or loss of repressive marks enrichment.

**Figure 7:**
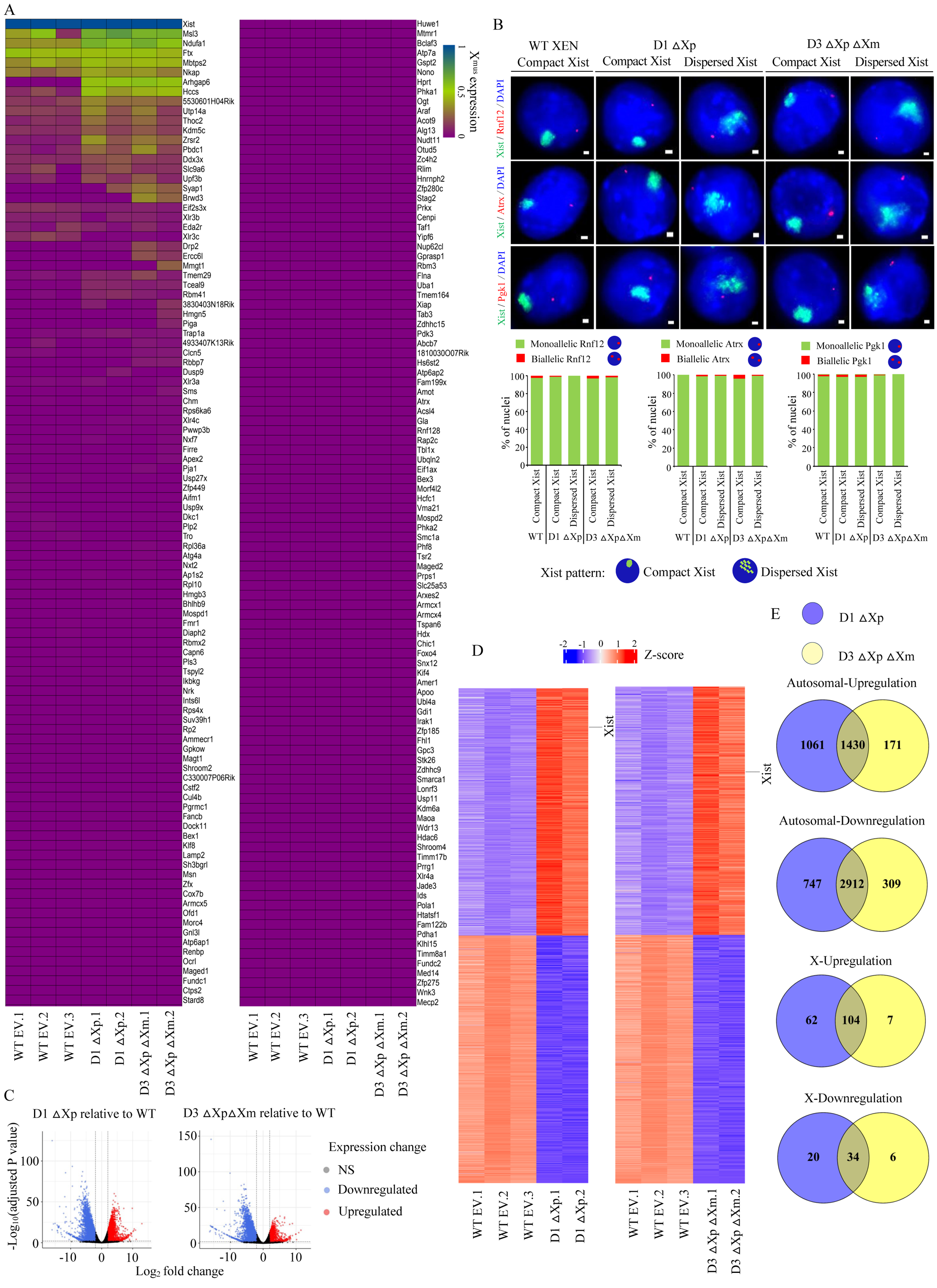
Xist upstream deletion affects autosomal gene expression but no effect on X-linked gene silencing. (A) Heat map showing the allelic expression pattern of different X-linked genes for D1 ΔXp.1, D1 ΔXp.2, D3 ΔXp ΔXm.1 and D3 ΔXp ΔXm.2 lines as compared to WT EV clones (WT EV.1, WT EV.2 and WT EV.3). (B) Representative images of RNA-FISH for Xist (green) and X-linked genes Rnf12, Atrx and Pgk1 (red) in WT, D1 and D3 deletion lines. Bottom: quantification of mono/biallelic expression of Rnf12 (n= 109, 109, 86, 116, 126), Atrx (n= 103, 85, 101, 159, 136) or Pgk1(n= 117, 118, 102, 120, 88) in nuclei having compact vs. dispersed Xist clouds. (C) Volcano plots representing the differentially expressed genes in WT vs D1 ΔXp and WT vs D3 ΔXp ΔXm lines. (D) Heat map showing the top 1000 dysregulated genes in the two deletions (D1 and D3) as compared to wildtype clones. (E) Venn-diagrams showing autosome and X-chromosome wise upregulated and downregulated genes for deletion D1 vs. deletion D3. Overlapping regions of the circles shows the commonality of the dysregulated genes between the two deletions.

**Figure 8.**
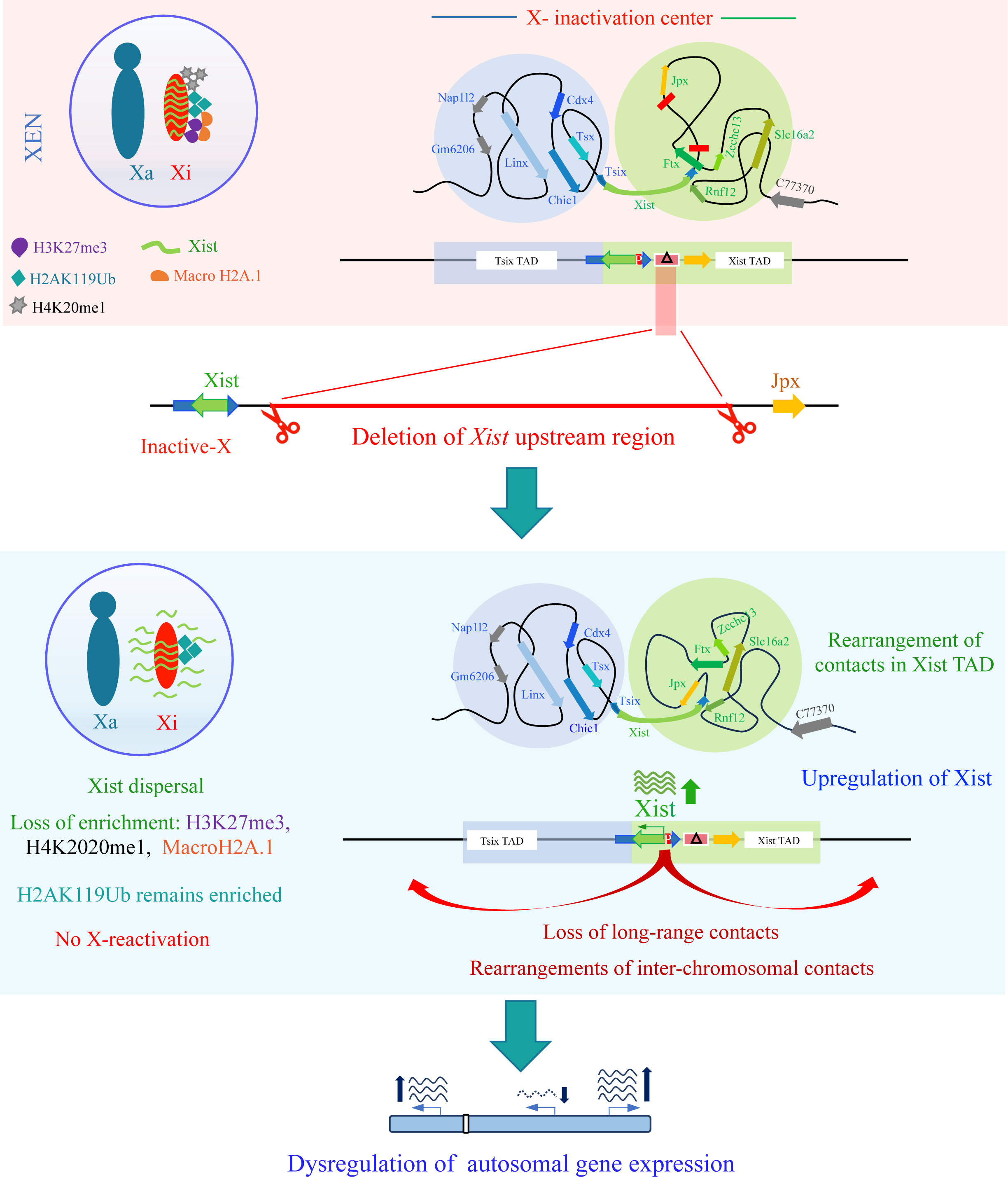
Graphical Summary. Xist upstream sequences are essential for maintaining proper TAD organization, Xist coating/expression and autosomal gene expression

Although there was not much effect on inactive-X gene silencing, we explored if there was any effect on autosomal gene expression because the expression of Xist and Tsix TAD related genes was affected in D1 ΔXp, and D3 ΔXpΔXm XEN cells. Interestingly, many autosomal genes showed differential expression in D1 ΔXp, and D3 ΔXpΔXm XEN cells compared to the WT EV clones (Fig. 7C, 7D). Moreover, many upregulated or downregulated genes were common between D1 ΔXp, and D3 ΔXpΔXm XEN cells (Fig. 7E). Collectively, we concluded that perturbation of inactive-X TAD and associated gene expression affects the autosomal gene expression.

## Discussion

Mammalian chromosomes are comprised of two major compartments: open chromatin and closed chromatin. Chromatin is further partitioned into TADs, whose boundaries are mediated through architectural proteins like CTCF and cohesion (Fudenberg et al., 2017; Krivega and Dean, 2017). It is believed that chromatin state and TADs play a collective role in fine-tuning gene expression (Fukuda et al., 2023; Kundu et al., 2017; Szabo et al., 2019). However, interplay between TADs and gene expression remains controversial (Cavalheiro et al., 2021; Tena and Santos-Pereira, 2021). Here, using murine XIC topology, we have addressed in what extent precise topological organization is important for gene regulation on the inactive-X chromosome in XEN cells.

The topological organization of the XIC has been studied in mouse cells undergoing random X-inactivation (Galupa, 2023; Galupa and Heard, 2018; Nora et al., 2012). It has been shown that the region of the XIC is partitioned into two antagonistically functioning TADs - Xist and Tsix-TADs (Fig. 1A). Based on 4C-seq analysis, we show that XIC in XEN cells with imprinted inactive-X have almost similar topological organization like cells with random X-inactivation (Fig 1). Furthermore, most of the CTCF binding sites at the Xist and the Tsix TADs in XEN cells were found to be very similar to the CTCF enrichment sites in NPC with random X-inactivation (Fig 1). Overall, the topological organization of the XIC in XEN cells having imprinted inactive-X is very similar to other cell types which undergo random X-inactivation. In converse, the enrichment of H3K4me1 and H3K27ac was different between XEN and differentiating ESC. On the other hand, we found few interactions of Xist locus with the neighbouring Tsix-TAD loci, suggesting evidence for inter-TAD interactions as reported previously (Fig 1C) (Despang et al., 2019; Galupa et al., 2020). However, inter-TAD contacts does not always mean to inter-TAD regulation (Despang et al., 2019).

Next, we explored the topological and transcriptional consequence upon deletion of *Xist* upstream sequences, which are a part of the Xist TAD and located near the boundary in XEN cells. Through chromosome conformation analysis, we found that deletion of Xist upstream sequence led to dramatic rearrangements of contacts within the TAD and loss of interaction outside the TAD (Fig 2). Since the alterations in the interactions were global and not limited to only the Xist TAD, it appeared that local deletion of a chromosomal segment could lead to disorganization of the entire chromosome and that TAD boundaries do not completely limit interactions with neighboring TADs as reported previously (Galupa et al., 2020). Interestingly, TAD alterations affected the expression of genes within Xist TAD as well as in neighbouring Tsix-TAD (Fig. 3). Specially, expression of Xist was significantly increased upon TAD alteration in Xist upstream deleted cells (Fig. 3).

On the other hand, deletion of the Xist upstream sequences (D1 deletion ∼6 kb) resulted in dispersal of Xist coating (Fig. 3). Investigation of these dispersed clouds showed multiple discrete Xist foci that spread along a large area of the nucleoplasm. It is highly possible that conformational changes of inactive-X upon deletion of *Xist* upstream sequence leads to the dispersal of Xist coating. Indeed, previous report has shown that spreading and coating of Xist RNA depend on the structure and the spatial configuration of the X-chromosome (Chen et al., 2016). Additionally, in Xist dispersed nuclei, there was loss of enrichment of different repressive chromatin marks from the inactive-X such as H3K27me3, H4K20me1 and MacroH2A.1 (Fig 5). Lack of enrichment of repressive histone modifications over an inactive-X that had dispersed Xist indicated that recruitment of such marks is dependent on the compact Xist coating. However, H2AK119ub was still strongly enriched on the inactive-X of dispersed Xist nuclei, suggesting that enrichment of H2AK119ub is not dependent of proper compact coating of Xist RNA (Fig 5). Our result is consistent with a recent study showing that during initiation of X-inactivation H2AK119ub is enriched even before Xist is accumulated compact manner on the inactive-X (Valledor et al., 2023). Segmental deletion of ∼3 kb (D3) close to *Jpx* recapitulated same phenotype related to Xist coating and loss of enrichment of repressive chromatin marks (Fig. 4). Interestingly, Xist dispersal and loss of repressive mark enrichment did not alter nucleolar association of the inactive-X (Fig. 6). Moreover, we found that Xist tethering proteins such as CELF1, CIZ1 were still robustly enriched in Xist dispersed nuclei indicating that enrichment of tethering proteins is not sufficient enough for proper Xist coating (Fig. 6). Surprisingly, despite of Xist dispersal and loss of enrichment of repressive chromatin marks, genes on the inactive-X remains silent. However, we observed dysregulation of autosomal gene expression though X-linked gene silencing maintained in these cells. This could be due to several reasons such as (1) upregulation of Xist and other TAD associated gene’s expression (2) binding of dispersed Xist RNA at other autosomal loci (3) change of contacts between X-chromosome and other autosomes leading to dysregulation of autosomal gene expression. Indeed, it was shown that deletion of X-chromosome mega-structure controlling element *Firre* and *Dxz4* did not affect X-linked gene silencing but led to dysregulation of many autosomal genes (Andergassen et al., 2019). Altogether, we conclude that *Xist* upstream sequences are required for maintaining proper TAD organization, Xist coating/expression and autosomal gene expression. Finally, our study provides significant insight and extends support into the role of proper genome organization in regulating transcriptional gene expression.

## Materials and Methods

### Data availability

Raw data for RNA-seq, ChIP-seq and 4C-seq generated in this study will be deposited to Gene Expression Omnibus (GEO) and made available to the public after publication. Upon request, data can be available by the corresponding author. The previously published dataset used for this study is available at GEO under the following accessions: ChIP-seq data for CTCF: GSE99991, H3K4me1 and H3K27ac in 24h differentiated ESCs (GSE116480) and Capture-C: GSE124596.

### Cell culture

XEN cells were cultured in Dulbecco’s modified eagle medium (DMEM) media (Himedia, #AL007A) as described previously (Naik et al., 2023). Media was supplemented with 20% fetal bovine serum (GIBCO, #10270-106), 3 mM L-Glutamine (GIBCO, #25030081), 1.5X MEM non-essential amino acids (GIBCO, #1140-050), Pen-Strep (GIBCO, #15140122), β-Mercaptoethanol (Sigma, #M3148). Cells were grown on gelatin (Himedia, # TCL059) coated plates at 37 °C with 5% CO2 and passaged using 0.05% Trypsin.

### CRISPR-based knockout cell line generation

Different *Xist-*upstream deletion clones (D1, D2 and D3) was generated through CRISPR/Cas9 based approaches using pairs of small guide RNAs (sgRNAs) targeting specific genomic regions, designed using an online tool CHOPCHOP (https://chopchop.cbu.uib.no) and checked against off-target effects using the tool Cas-offinder (http://www.rgenome.net/cas-offinder/). sgRNAs used for different deletion clones is as follows: (1) For D1-sgRNA1: CAAGATTACTATTTTACCCCAGG; sgRNA2: GACCAAGGCGGGGTAGAAGAAGG. (2) For D2-sgRNA1: CAAGATTACTATTTTACCCCAGG; sgRNA2: AATTATAGGAG**G**GCTTCACATGG. (3) For D3-sgRNA1: AATTATAGGAG**G**GCTTCACATGG; sgRNA2: GACCAAGGCGGGGTAGAAGAAGG. In brief, sgRNAs were cloned individually into the plasmid <u>pSpCas9(BB)-2A-Puro (PX459)</u> <u>V2.0</u> (Addgene, #62988) and then transfected into XEN cells using Xfect^TM^ transfection reagent (Clontech, #631318) in presence of 1X Opti-MEM (GIBCO, #31985-070). Next, after selection through Puromycin (3ug/ml), clones were FACs sorted in 96-well plates as single-cell and expanded for PCR screening followed by Sanger-sequencing. The nature of the heterozygous clones (carrying paternal or maternal deletion) was differentiated through PCR conducted using primers spanning single-nucleotide polymorphisms. All primers used for the screening is listed in Table S1.

### RNA FISH

For RNA FISH, we generated double-stranded RNA-FISH (dsRNA FISH) probes through random priming of BAC templates (Xist, Atrx, Rnf12, Pgk1) using the Bioprime labelling kit (Invitrogen, #18094-011). Probes were labelled using Cy3-dUTP (Enzo Life Sciences, #ENZ-42501) or Cy5-dUTP and purified through ProbeQuant G-50 Micro columns (Cytiva, #28903408). Strand-specific RNA-FISH (ssRNA FISH) probes were produced using the MAXIscript T3/T7 kit (Ambion, #AM1326). ssRNA FISH probes were labelled using FITC-UTP and purified through Mini Quick Spin RNA columns (Roche, #11814427001). Labelled double-stranded and ssRNA FISH probes were then precipitated in presence of 0.3M sodium acetate (Sigma, #71196) and 0.5M ammonium acetate (Sigma, #09691) respectively along with 300 μg of Yeast tRNA (Invitrogen, #15401011), 150 μg of sheared Salmon sperm DNA (Invitrogen, #15632-011) and absolute ethanol (Hayman, #F205220) at 13,000 rpm for 20 mins at 4°C. The pellet was washed with 70% and then 100% ethanol, vacuum-dried and resuspended in deionized formamide (VWR Life Sciences, #0606) and then denatured at 95°C for 10 mins followed by immediate snap-freezing on ice. A hybridization solution containing 20% Dextran sulphate (SRL, #76203), 2X SSC (SRL, #12590) was mixed with the denatured solution and the probes were then stored at -20°C until the use.

For RNA-FISH, cells mounted on coverslips or on which IF had been conducted (fixed after IF with 2% PFA), were first dehydrated through an ethanol series of 70%, 85%, 95% and 100% for 2 mins each and subsequently air-dried. Cells were then hybridised with double-stranded or single-stranded probes for overnight at 37°C. Next day, cells were washed thrice with 2X SSC/50% Formamide, 2X SSC, and twice with 1X SSC for 7 mins each at 37°C. DAPI (Invitrogen, #D1306) diluted to 1:1000 was added at the third 2X SSC wash. The coverslips were finally mounted using Vectashield (Vector Labs, #H1000) and sealed with nailpolish before visualisation and image acquisition.

### Immunofluorescence(IF) and followed by RNA-FISH

For Immuno-fluorescence, XEN cells were grown on gelatin (Himedia, #TCL059)-coated coverslips, followed by their permeabilization with ice-cold Cytoskeletal buffer (composed of 300 mM Sucrose, 100mM NaCl, 3mM MgCl2 and 10mM PIPES buffer, pH 6.8) for 30 s, followed by treatment with CSK buffer containing 0.4% Triton X-100 for 30 s and another round of CSK buffer for 30 s. Cells were then fixed using 3% paraformaldehyde (Electron Microscopy Sciences, #15710) for 10 min, washed 3 × with 70% ethanol and then stored at - 20 °C in 70% ethanol before conducting IF or RNA-FISH. For IF, XEN cells plated on 22μm coverslips (Blue Star) were firstly washed 3× with PBS (Himedia, #TL1006) for 3 mins while shaking, following which they were incubated in blocking buffer composed of 0.5 mg/ml BSA (New England Biolabs, #B9001S), 80 units of RNAse-OUT RNAse inhibitor (Invitrogen, #15401-029), 0.2% Tween-20 (Promega, #H5152) and PBS for 30 mins in a humid chamber humidified using PBS/0.2% Tween-20 at 37°C. Cell samples were then incubated with appropriately diluted primary antibody for 1 hr at 37°C. The anti-trimethyl Histone H3 (Lys27) antibody (Merck, #07-449) was used at a dilution of 1:2500, the anti-Ubiquityl-Histone H2A (Lys119) antibody (CST, #8240T) at a dilution of 1:1600, the H4K20me1 antibody (ABclonal, # A2370) at a dilution of 1:80, the MacroH2A.1 antibody (ABclonal, #A9059) at a dilution of 1:400, the CIZ1 antibody (ABclonal, #17349) at a dilution of 1:500, the CUGBP1 antibody (Invitrogen, #MA1-16675) at a dilution of 1:80 and the FBL antibody (ABclonal, #A1136) at a dilution of 1:200. Samples were then washed using PBS/0.2% Tween-20, 3× for 3 minutes each, while shaking at 37°C following which they were incubated with 1:300 diluted fluorescently-conjugated secondary antibody (Alexa-Fluor, Invitrogen). Cell samples were then washed 3× with PBS/0.2% Tween-20 for 3 minutes each at 37°C and processed further for RNA-FISH as described above.

### RT-qPCR

For quantitative real-time PCR, RNA was isolated from XEN cells following Trizol (Ambion,) method. RNA was first reverse-transcribed using Primescript 1^st^ strand cDNA Synthesis kit (TaKaRa, #6110A) with random hexamers at 30°C for 10 mins, 42°C for 60 mins followed by heat inactivation of the reverse transcriptase enzyme at 95°C for 5 mins. Real time PCR was performed on the cDNA using KAPA SYBR FAST qPCR Master Mix (2X) kit (Roche, #KK4618) with intron spanning primers in a QuantStudio 3 Real-time PCR system. All Ct values were normalized against TBP. A 2^-ΔΔCt method was followed to calculate all relative fold changes of the genes. All RT-qPCR primers used for this study have been listed in Table S2.

### RNA-sequencing and analysis

RNA-sequencing and analysis was performed as described previously with few modifications (Naik et al., 2023). In brief, we isolated total RNA from XEN cells using TRIzol (Life technologies #15596-026) following the manufacturer protocol. Next, we checked quality of the RNA through agarose-gel electrophoresis followed by Bioanalyzer. RNA samples, which showed RIN value above 7, was used for the library preparation. RNA-Seq library was prepared following the manufacturer’s protocol using the NEBNext/truseq library preparation kit for Illumina. After successful library preparation, we sequenced libraries on Illumina platform using pair-end (2 X 150 bp) chemistry. For data analysis, we first performed FastQC of the transcriptomic sequencing reads and adapter content were removed using Trim Galore (v0.6.6). Further, we removed the rRNA readss using RiboDetector (v0.2.7) (Deng et al., 2022). Next, we used STAR (2.7.9a) to map the transcriptomic sequencing reads to the mouse genome GRCm38 (mm10) (Dobin et al., 2013). Counting of reads was done using featureCounts (2.0.1) (Liao et al., 2014). For differential gene expression analysis, we used R package DESeq2 tool (Love et al., 2014).

### Allele-specific RNA-seq analysis

We performed allele-specific analysis of RNA-seq data following the previously described method (Naik et al., 2023, 2021, 2022). In brief, first we constructed in silico reference genome through incorporation of strain-specific SNPs in to the mm10 reference genome using VCF tools. We accessed strain-specific SNPs from Mouse Genomes Project (https://www.sanger.ac.uk/science/data/mouse-genomes-project). Next, transcriptomic reads were mapped separately to parental genomes using STAR. We only considered those SNPs, which had minimum read counts of 10 per SNP site. We considered only those genes which had at least 2 informative SNPs for our analysis. We calculated an average of SNP-wise reads to estimate allelic read counts. We calculated allelic ratio using the following formula: (Allele-Mus or Allele-Mol/ Allele-Mus + Allele-Mol).

### ChIP-sequencing

For Chromatin immunoprecipitation (ChIP), we used the Simple ChIP Enzymatic Chromatin IP Kit with Magnetic beads (Cell Signalling Technology #9003) and followed the manufacturer’s instructions. In brief, cells were fixed with 1% formaldehyde (Sigma #F8775) in DMEM at room temperature for 20 min and then quenching was done with 125mM glycine for 5min at room temperature. Cells were then washed with ice-cold PBS and then mechanically harvested using a cell scraper. Cells were lysed on ice with the lysis buffer provided in the kit and nuclei were isolated. Chromatin was fragmented by partial digestion with 0.5 μl micrococcal nuclease per IP followed by sonication with Biorupter pico (Diagenode) for 20 cycles 30 sec on/off. Next, the cross-linked chromatin was immunoprecipitated with ∼1μg of CTCF antibody (Diagenode), H3K4me1 antibody (Diagenode), H3K27ac antibody (Diagenode) and isotype control IgG (provided in the kit) separately from the sample. 2% input chromatin DNA were kept aside before IP which acts as a control to normalize signal from ChIP enrichment. The immunoprecipitated chromatin and input chromatin were reverse cross-linked and subjected purification. Purified ChIP DNA was quantified using Qubit. After initial quality control, ChIP DNA were subjected to library preparation. ChIP library was sequenced on Illumina Hiseq platform using 150×2 paired end chemistry.

### Allele-specific ChIP-seq analysis

Allele-specific ChIP-seq analysis was performed following the methods as described previously (Parichitran et al., 2023). In short, first, we created a GRCm38 (mm10) N-masked genome in-silico using SNPsplit genome preparation. Next, we used Bowtie2 to map the ChIP-seq reads to the in-silico created N-masked genome (B and SL, 2012). Blacklisted regions were removed from our analysis according to the encode consortium. SNPsplit (0.4.0) was then used to create allele-specific BAM files by segregating the aligned reads into two distinct alleles (F and SR, 2016). Enrichment ratio log2(ChIP/input) was calculated using bamCompare function using deepTools (v.3.5.0) with parameter centerReads, ignoreDuplicates, 20-bp bin, smooth length 200 bp, BPM normalization (Ramírez et al., 2016).

### Circular Chromosome Conformation Capture (4C) sequencing and analysis

XEN cells (∼10 million) were grown under appropriate conditions and fixed using 1.5% formaldehyde (Sigma, #F8775) while shaking at 40 rpm. Formaldehyde was quenched using 125mM Glycine (Cell Signalling technology, #7005). Cells were washed with ice-cold PBS (2X), scraped and pelleted before storing them at -80°C. Cells were lysed using Tris-Cl pH 8.0 (10 mM), NaCl (10 mM), NP-40 (0.2%), PIC (1X), homogenised by a dounce homogenizer (15 strokes with each of pestle A and pestle B) and pelleted down. The nuclei pellet was then washed 2X with ice-cold 1X DPBS and resuspended in restriction digestion buffer. The pellet was then incubated at 65°C in presence of 1% SDS to get rid of uncrosslinked proteins for 8 mins and then immediately transferred to ice. The SDS was neutralised using 10% TritonX-100, following which the chromatin was digested using 100 units of the 1^st^ cutter NlaIII (NEB, #R0125L) for overnight at 37°C. The digested chromatin was then subjected to ligation using T4 DNA ligase (Invitrogen, #) and T4 DNA Ligase buffer (NEB, #B0202S) at 16°C for 4 hours. The ligated samples were decrosslinked overnight using Proteinase K (Roche, #3115887001), PCR purified and ethanol precipitated. 50 units of DpnII (NEB, #R0543M) was used for the second restriction digestion following which samples were again ligated, purified and precipitated similarly to obtain the 4C library. The 4C library was treated with RNAse A, and purified using the QIAquick PCR purification kit. The concentration of the 4C library was quantified and PCR was conducted using specific viewpoint oligos. The oligo sequences used for 4C are summarised in Table S3. The amplified product was then PCR purified and subjected to next-generation sequencing using HiSeq 2500. Data were analyzed using R tool pipe4c (v1.16) (https://github.com/deLaatLab/pipe4C) pipeline with default parameters (Krijger et al., 2020). In brief, reads are demultiplexed followed by mapped using bowtie2 against BSgenome.Mmusculus. UCSC.mm10. For Integrative Genomics Viewer (IGV) visualization normalized and smoothened bigwig files were used.

### Data visualization and plots

Plots were generated through R version 4.2.1 using ggplot2.

### Microscopy

RNA-FISH and IF images were captured using Olympus IX73 Research Inverted microscope and cellSens [Ver.2.1] Life Science imaging software equipped with a 100X oil-immersion objective. Z-stacks for each image were acquired at a 0.25 μm step size and EFI processing was conducted to get the maximum intensity projection of the image. For area and sphericity measurements, Xist clouds were defined in the wild-type and mutant cells using the ROI (region of interest) option and the measurements were derived from the measurements and ROI tool of the cellSens imaging software. Intensity profile of H3K27me3 across the *Xist* cloud was measured using the line Profile tool window and measurements were exported as a chart for plotting the profile.

## Acknowledgement

We thank Prof. Sundeep Kalantry, University of Michigan for providing BACs and cell line. This study is supported by DBT grant (BT/PR30399/BRB/10/1746/2018), DST-SERB (CRG/2019/003067), DBT-Ramalingaswamy fellowship (BT/RLF/Re-entry/05/2016) and Infosys Young Investigator grant award to SG. SM and Avinchal acknowledge University of Grant Commision (UGC) for the fellowship. RB would like to acknowledge council of scientific and industrial research (CSIR), India for the fellowship. LSB acknowledge DST Women scientist award. We thank NCBS genomic core facility for the sequencing

## Authors Contribution

SG supervised and acquired the funding for the study. SG and SM conceptualized the study. SM, LSB, RB, Avinchal, AK and GSB performed experiments. HCN and NS performed bioinformatic analysis. DN supervised and helped with the 4C experiments. SM wrote the first draft of the manuscript. HCN and LSB contributed to writing the manuscript. SG and SM edited and proofread the manuscript. The final manuscript was approved by all the authors.

