## Supplementary figures for "Deletion of *Xist* upstream sequences alters TAD interactions and leads to defects in Xist coating and expression"

Fig. S1

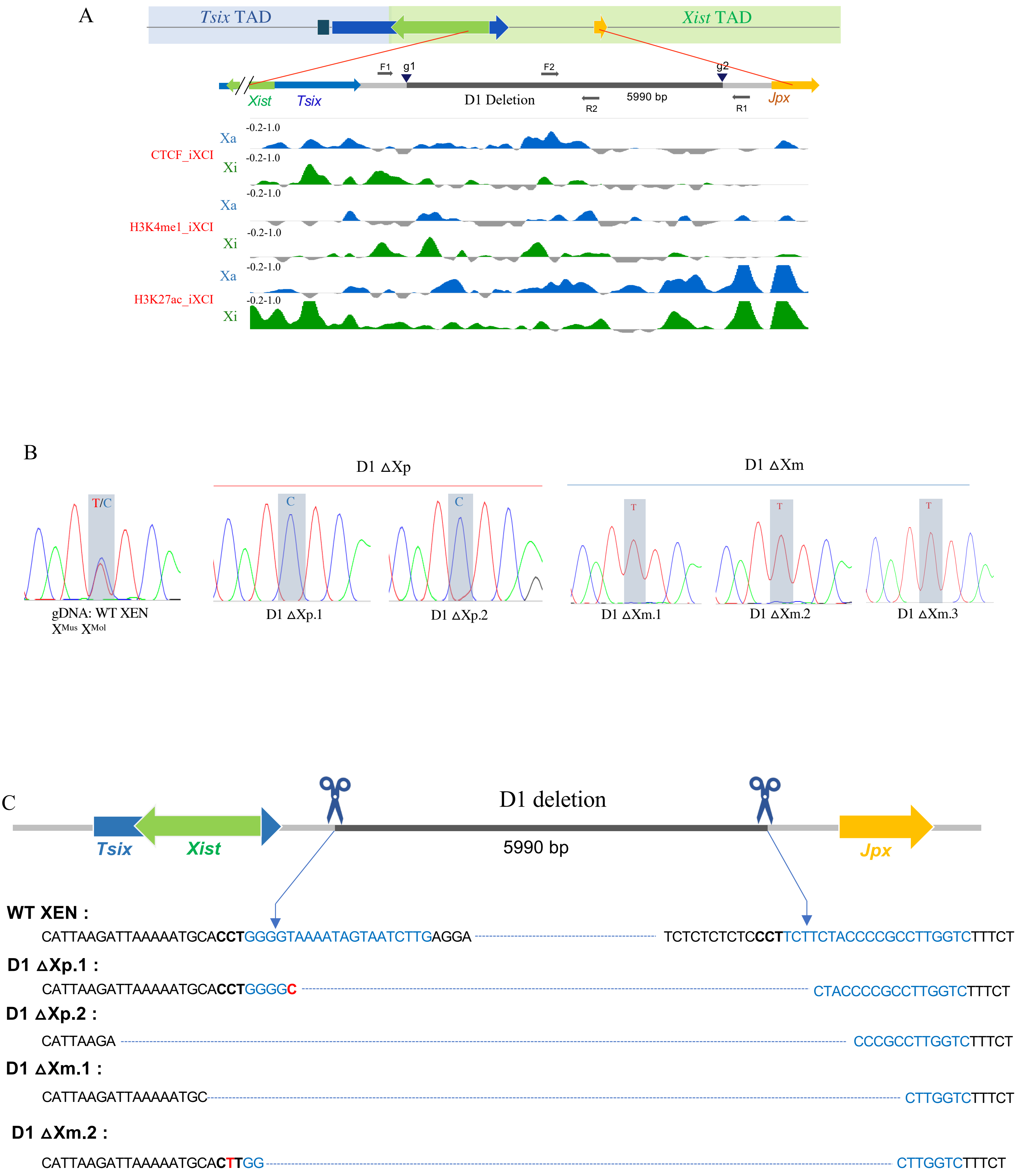


Blue sequences indicate gRNAs, Black sequences are gRNA flanking regions, **Black and bold nucleotides indicate PAM**

**Fig. S1, related to Figure 2:** (A) ChIP-seq enrichment of CTCF, H3K4me1 and H3K27ac in active (blue) and inactive-X (green) at the Xist upstream region targeted for deletion in WT XEN cells. (B) Representative Sanger-sequencing chromatogram of genomic DNA from WT and Xist-upstream deleted clones. SNP analysis showing the heterozygous deletions (either paternal or maternal) in different D1 deleted clones. (C) Schematic showing PCR-based analysis of the deleted allele for all D1 deleted clonal lines. The PAM sequences are indicated as bold black characters. The Cas9 recognition sequences are indicated in blue. Indels and substitutions are highlighted in red. Dotted lines for the deletion clones indicate the deleted region.

Fig. S2

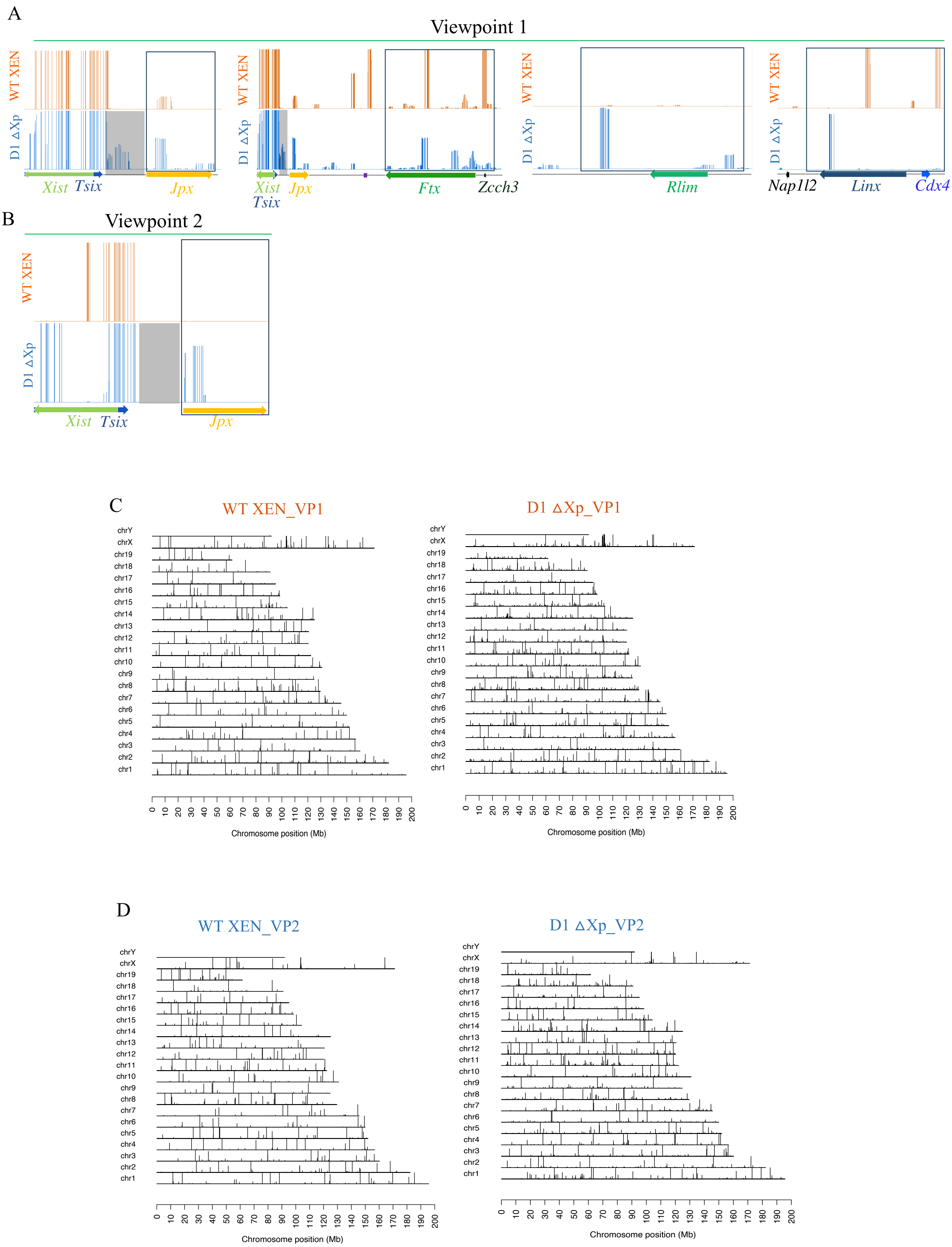
A
Viewpoint 1
WT XEN
D1 ΔXp
Xist
Tsix
Jpx
Ftx
Zcch3
Rlim
Nap1l2
Linx
Cdx4
B
Viewpoint 2
WT XEN
D1 ΔXp
Xist
Tsix
Jpx
C
WT XEN_VP1
D1 ΔXp_VP1
chrY
chrX
chr19
chr18
chr17
chr16
chr15
chr14
chr13
chr12
chr11
chr10
chr9
chr8
chr7
chr6
chr5
chr4
chr3
chr2
chr1
0 10 20 30 40 50 60 70 80 90 100 110 120 130 140 150 160 170 180 190 200
Chromosome position (Mb)
D
WT XEN_VP2
D1 ΔXp_VP2
Chromosome position (Mb)

**Fig. S2, related to Figure 2:** (A) Representation of the differential 4C interactions between the WT and D1 ΔXp lines in the Xist and the Tsix TAD with respect to Xist VP 1. The deleted Xist upstream region have been shaded in grey. (B) The differential 4C interactions between the WT and the D1 ΔXp line for Xist viewpoint 2 have been highlighted for the *Jpx* region within a box. The D1 deleted region have been shaded in grey. (C) and (D) The normalised genome-wide interactions for Xist viewpoint 1 (VP1) and Xist viewpoint 2 (VP2) respectively, across all mouse chromosomes in WT and D1 ΔXp line are represented.

Fig. S3

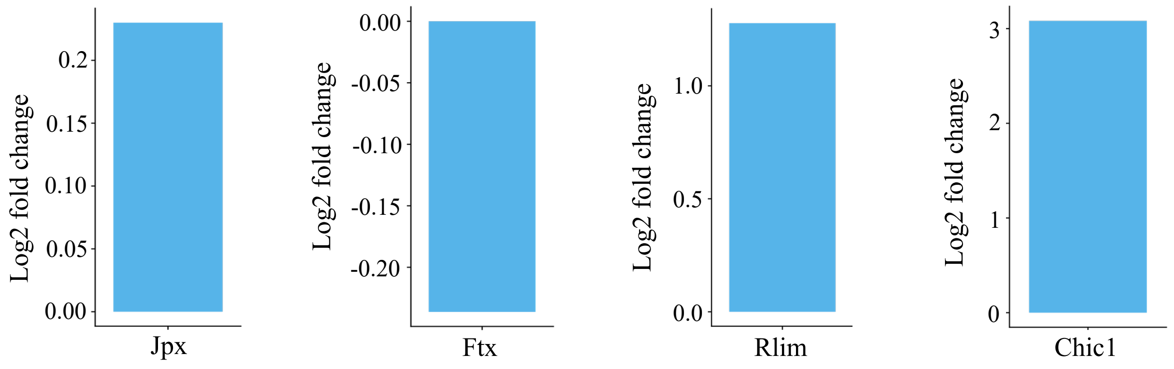


D1 ΔXp over WT EV

**Fig. S3, related to Fig. 3:** $Log_2$ fold change in expression levels of *Jpx*, *Ftx*, *Rnf12* and *Chic1* in D1 ΔXp lines over WT EV as observed through RNA-sequencing.

Fig. S4

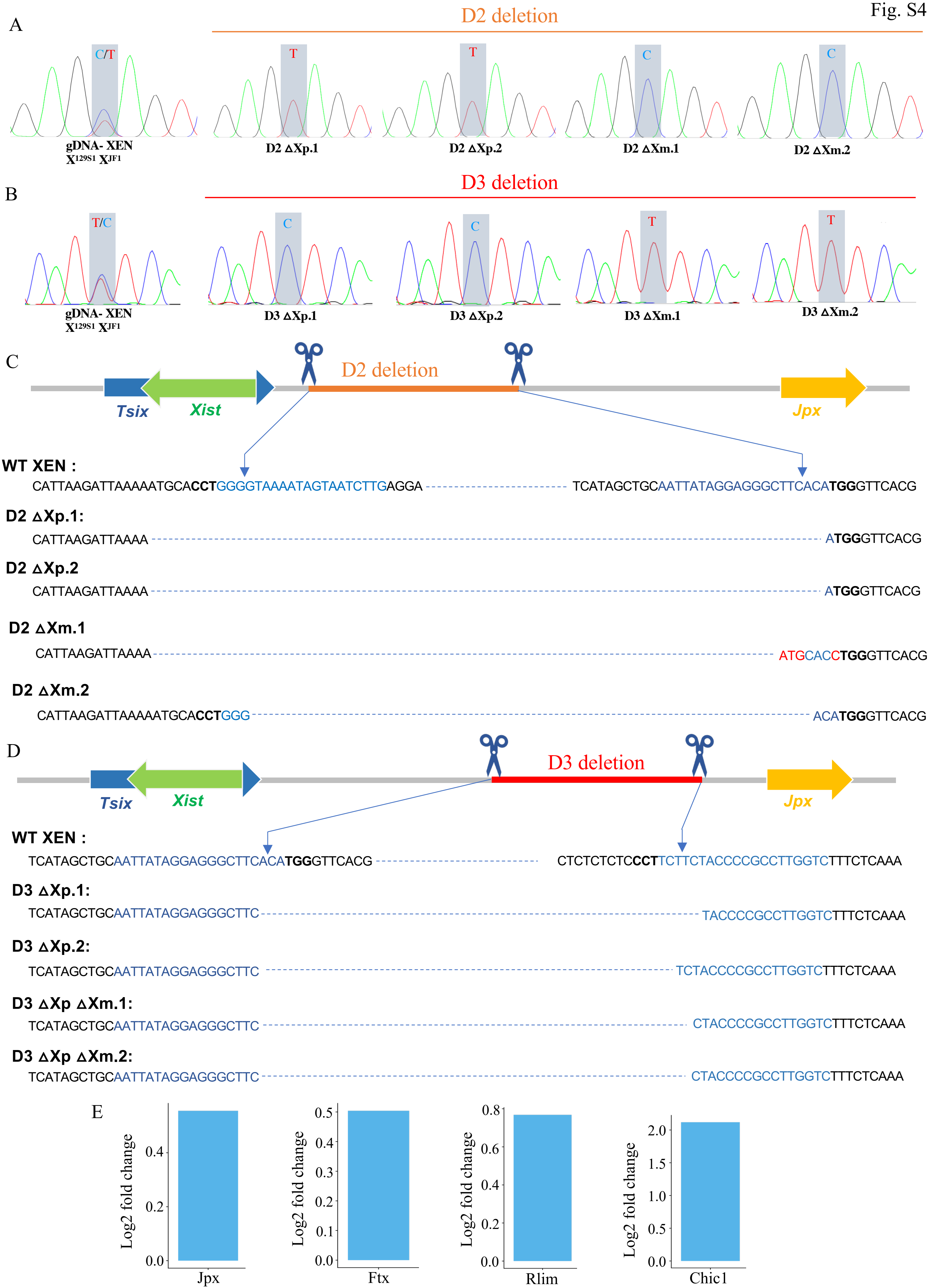

A
D2 deletion
C/T
T
T
C
C
gDNA- XEN
X129S1 XJF1
D2 ΔXp.1
D2 ΔXp.2
D2 ΔXm.1
D2 ΔXm.2
B
D3 deletion
T/C
C
C
T
T
gDNA- XEN
X129S1 XJF1
D3 ΔXp.1
D3 ΔXp.2
D3 ΔXm.1
D3 ΔXm.2
C
D2 deletion
Tsix
Xist
Jpx
WT XEN :
CATTAAGATTAAAAATGCACCTGGGGTAAAATAGTAATCTTGAGGA
TCATAGCTGCAATTATAGGAGGGCTTCACATGGGTTCACG
D2 ΔXp.1:
CATTAAGATTAAAA
ATGGGTTCACG
D2 ΔXp.2
CATTAAGATTAAAA
ATGGGTTCACG
D2 ΔXm.1
CATTAAGATTAAAA
ATGCACCTGGGTTCACG
D2 ΔXm.2
CATTAAGATTAAAAATGCACCTGGG
ACATGGGTTCACG
D
D3 deletion
Tsix
Xist
Jpx
WT XEN :
TCATAGCTGCAATTATAGGAGGGCTTCACATGGGTTCACG
CTCTCTCTCCCTTCTTCTACCCCGCCTTGGTCTTTCTCAAA
D3 ΔXp.1:
TCATAGCTGCAATTATAGGAGGGCTTC
TACCCCGCCTTGGTCTTTCTCAAA
D3 ΔXp.2:
TCATAGCTGCAATTATAGGAGGGCTTC
TCTACCCCGCCTTGGTCTTTCTCAAA
D3 ΔXp ΔXm.1:
TCATAGCTGCAATTATAGGAGGGCTTC
CTACCCCGCCTTGGTCTTTCTCAAA
D3 ΔXp ΔXm.2:
TCATAGCTGCAATTATAGGAGGGCTTC
CTACCCCGCCTTGGTCTTTCTCAAA
E
Log2 fold change
0.0
0.2
0.4
Jpx
Log2 fold change
0.0
0.1
0.2
0.3
0.4
0.5
Ftx
Log2 fold change
0.0
0.2
0.4
0.6
0.8
Rlim
Log2 fold change
0.0
0.5
1.0
1.5
2.0
Chic1

**Supplementary Fig.S4, related to Fig. 4:** (A) and (B) SNP based Sanger-sequencing analysis of the undeleted allele of each clone to differentiate between the paternal and the maternal deletions. (A) For the D2 deletion, "C" represents the paternal/*Mus* allele and "T" the maternal/*Mol* allele. Presence of T at the polymorphic site indicates wildtype allele of the clone is maternally derived and hence the paternal allele is deleted for the D2 region. (B) For the D3 deletion, T" indicates the paternal/*Mus* allele and "C" indicates the maternal/*Mol* allele. (C) and (D) PCR based evaluation of the deleted region in D2 and D3 deletion lines respectively. Black bold characters highlight the PAM sequences, blue characters highlight gRNA sequences. The blue dotted lines for the deleted clones specify the deleted region. Indels and substitutions are highlighted as red characters. (E) $Log_2$ fold change in expression levels of *Jpx*, *Ftx*, *Rnf12* and *Chic1* in D3 ΔXp ΔXm lines (n=2) over WT EV (n=3) as obtained from RNA-seq analysis.

### Table S1: Primer sequences

| **Experiment** | **Primer Name** | **Sequence** |
|---|---|---|
| PCR D1 del | F1 | GAAGTTAGAAATCCACCTGCCTCC |
| PCR D1 del | R1 | CCTCATCACCCCTTTTTGTCCA |
| PCR D1 and D3 wt and snp | F2 | GATGGCCTACTTGGCTTCACA |
| PCR D1 and D3 wt and snp | R2 | ATCTTGACTGGCCATGGTAGC |
| PCR D2 del | F3 | GGGGACTCTTCACTTTCTATTACT |
| PCR D2 del | R3 | CCATGAAGACAATAGACTTTAACCC |
| PCR D2 wt | F4 | CAGCATGGAGGCTGAACGAT |
| PCR D2 wt | R4 | GGTCACGGAACCAACTCCTG |
| PCR D3 del | F5 | CCTTGTTTACTGAGGCAAGATC |
| PCR D3 del | R5 | CTCAGGCCTTTGGCAATAGAC |

### Table S2: RT-qPCR primers:

| Primer name | Sequence |
|---|---|
| Xist 9574 FP ex 2 | CAAGAAGAAGGATTGCCTGGA TTT |
| Xist 9720 RP ex 3 | GCGAGGACTTGAAGAGAAGTTCTG |
| Jpx qFP | GCACCACCAGGCTTCTGTAAC |
| Jpx qRP | GGATTAATGATCAGGGCATGT |
| Ftx qFP | GCCGCTTTTGAAGAGATAAACT |
| Ftx qRP | TGATTATCCTCATGTGTTGCTG |
| Rnf12 qFP | GGTCCACCACCACAGAGC |
| Rnf12 qRP | TGACCACTTCTTGTTGTATTTCC |

### Table S3: 4C oligos

| Viewpoint | Forward Primer | Reverse Primer |
|---|---|---|
| 1 | **VP1 WT FP:** AAT GAT ACG GCG ACC ACC GAA CAC TCT TTC CCT ACA CGA CGC TCT TCC GAT CT GACCGAGGCCCCGTTGGGCATG<br><br>**VP1 Mut FP:** AAT GAT ACG GCG ACC ACC GAA CAC TCT TTC CCT ACA CGA CGC TCT TCC GAT CT TGCCGAGGCCCCGTTGGGCATG | **VP1 RP:** CAAGCAGAAGACGGCATACGA GCCTCTTATTTGCGTGTACTGTTG |
| 2 | **VP2 WT FP:** AAT GAT ACG GCG ACC ACC GAA CAC TCT TTC CCT ACA CGA CGC TCT TCC GAT CT GA TACGTAAGCTCAGTGACATG<br><br>**VP2 Mut FP:** AAT GAT ACG GCG ACC ACC GAA CAC TCT TTC CCT ACA CGA CGC TCT TCC GAT CT TG TACGTAAGCTCAGTGACATG | **VP2 RP:** CAAGCAGAAGACGGCATACGACT TCTGGTCTCTCCGCCTTCAGC |
